# Creating bacterial genomic diversity through large-scale reconfigurations reveals phenotype robustness to organizational genome change

**DOI:** 10.64898/2026.01.06.698047

**Authors:** M. Victoria Barja, Andrey G. Gomes de Oliveira, Leticia Larotonda, Lucia Vigezzi, Julanie Rogers, Ian T. Paulsen, Alfonso Soler-Bistué, Briardo Llorente

## Abstract

The ability to generate genomic diversity expands opportunities for understanding and engineering biology. Here, we demonstrate on-demand generation of diversity in bacterial genome configurations and its application to probing physiology under altered genome organization. We engineered the fast-growing bacterium *Vibrio natriegens* to enable large-scale stochastic duplications, translocations, inversions, and deletions, producing populations with a wide range of genome arrangements. We investigated phenotypic robustness to genome reconfigurations and found that distinct genome organizations can support stable physiology, indicating that bacteria may tolerate chromosomal alterations more readily than previously appreciated. Our work provides a framework for advancing the understanding and engineering of bacterial genomes and suggests that genome reconfigurations may contribute to phenogenetic drift, allowing for evolutionary exploration while preserving phenotype.

## Introduction

Elucidating the associations between genome organization and phenotype has significant implications for understanding and engineering life. Genome rearrangement is a universal process in bacteria with profound effects on phenotype and evolution (*1-7*), potentially altering in a single step the level of expression of multiple genes, their order, their number, and genome interactions within chromosomal regions and with cell components.

Previous studies have systematically deleted, inverted, and relocated bacterial chromosome sections (*8-13*). While these approaches are powerful, a more disruptive methodology allowing genome manipulation on an unprecedented scale, Synthetic Chromosome Recombination and Modification by LoxP-mediated Evolution (SCRaMbLE), was developed as part of the synthetic yeast genome project Sc2.0 (*14*). The SCRaMbLE method relies on the genomic integration of multiple symmetrical *loxP* (*loxPsym*) recombination sites that are substrates for the Cre recombinase. Induction of Cre activity drives concurrent stochastic rearrangements that can eliminate, duplicate, invert, and translocate chromosome segments flanked by *loxPsym* sites, allowing the exploration of a diverse space of genome variants (*15, 16*). To date, SCRaMbLE has been successfully applied in *Saccharomyces cerevisiae* yeast and, more recently, in human cells (*17*), but its implementation to generate genome reconfigurations in bacteria has remained unexplored until now.

In this study, we engineered the SCRaMbLE system into the emerging model and industrial workhorse bacterium *Vibrio natriegens*, the fastest-dividing organism known, offering unique advantages for accelerated experimentation. We demonstrate that diverse large-scale genome reconfigurations can be produced within *V. natriegens* populations. We also assessed phenotypic robustness to genome changes, identifying multiple strains with distinct reconfigured genomes that maintained, or even improved, growth rates compared to the wild-type *V. natriegens* strain.

## Results

### Engineering *Vibrio natriegens* for enabling large-scale genome reconfigurations

The genome of *V. natriegens* comprises two circular chromosomes, a primary 3.25 Mbp chromosome (chromosome 1) and a 1.92 Mbp secondary chromosome (chromosome 2). Nearly all postulated essential genes (272 of 278 genes) are encoded on chromosome 1 (*18*). Chromosome 2, although conserved across the *Vibrio* genus, is thought to primarily encode nonessential functions and serve as an evolutionary test bed for horizontally acquired genes (*19, 20*).

To construct a *V. natriegens* strain capable of undergoing diverse large-scale genome reconfigurations, we engineered chromosome 1 based on its many genomic elements of interest and greater genetic potential for future studies. To minimize unintended cell alterations and preserve a near-wild-type phenotype and fitness, we defined four design principles for *loxPsym* site integration.

First, *loxPsym* integrations were targeted to intergenic regions between open reading frames (ORFs) in opposite orientations to avoid polar effects that could alter the expression of downstream genes (*21*). Second, we avoided regions near the chromosome replication origin (*oriC*) to prevent potential cell cycle disruptions (*22, 23*). Third, we excluded intergenic regions adjacent to essential genes to minimize the risk of detrimental effects. Finally, a lesson learned from the synthetic yeast genome project Sc2.0 is that the SCRaMbLE system leads to deletions occurring with a relatively high incidence (*14, 24*). To reduce the occurrence of deletions and favor large-scale duplications, inversions, and translocations, we defined a fourth design principle consisting of integrating *loxPsym* sites interspacing relatively large chromosome segments containing annotated essential genes. The rationale behind this fourth design principle is to preferentially lead to duplications, inversions, and translocations by increasing the essential genetic content of each segment to favor their retention in reconfigured genomes, as deletions would eliminate essential genes and cause lethality. The final design comprised 14 *loxPsym* integrations distributed across chromosome 1, partitioning it into 14 segments ranging from 0.15 to 0.52 Mbp, with the chromosome *oriC* and terminus (*ter*) located in segments 14 and 7, respectively (Figure 1A).

**Figure 1.**
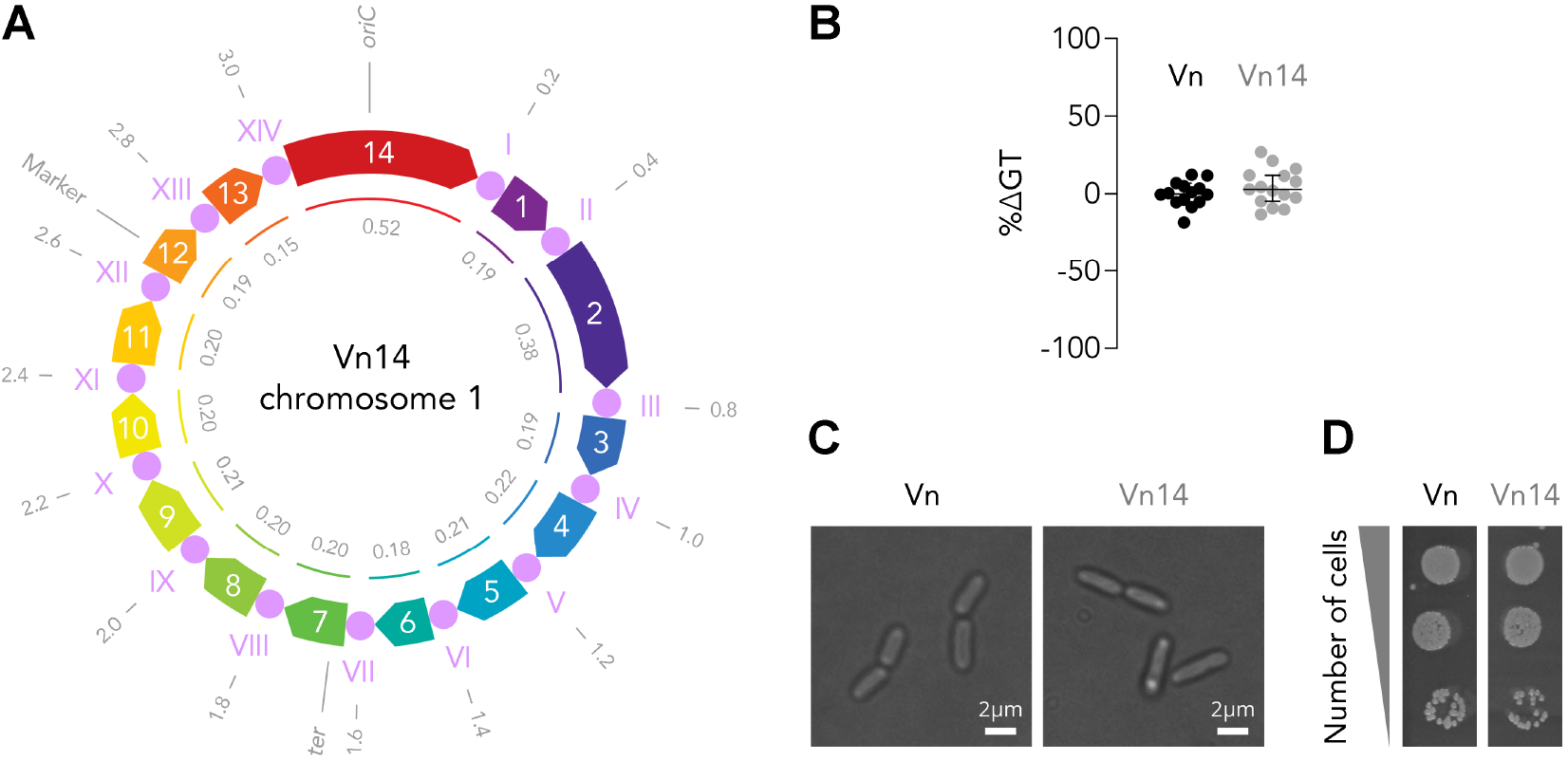
Design and profiling of *Vibrio natriegens* engineered for large-scale genome reconfigurations. (**A**) Engineering design of *V. natriegens* strain Vn14 chromosome 1. The outer circle indicates the genomic positions (Mbp) of *loxPsym* integrations, the chromosome replication origin (*oriC*), terminus (*ter*), and selection marker. The central circle shows the 14 *loxPsym* integrations (I to XIV) interspacing 14 chromosome segments represented as pentagons (1 to 14). Each chromosome segment is uniquely colored, and the direction of the pentagon represents the orientation. The inner circle indicates the size (Mbp) of each chromosomal segment. (**B**) Growth comparison of wild-type *V. natriegens* (Vn) and strain Vn14. Generation time (GT) results are presented as percentage of variation (%ΔGT) of the median with 95% CI relative to Vn (n = 12). (**C**) Visualization of Vn and Vn14 bacteria. (**D**) Fitness testing of Vn and Vn14 on LB3 agar. Serial 10-fold dilutions at 24 h.

We integrated the 14 *loxPsym* sites into the specified positions of chromosome 1 through multiple rounds of Multiplex Genome Editing by Natural Transformation (MuGENT) that alternatively swapped chloramphenicol (Cm^R^) and spectinomycin (Spec^R^) markers at the neutral *dns* locus (*25*). These markers were flanked by directional *loxP* sites - distinct from *loxPsym* - to enable Cre-mediated marker excision. Correct integration at each target site was verified by polymerase chain reaction (PCR) genotyping with primers flanking *loxPsym* insertions and by whole-genome sequencing (Figure S1 and Data File S1). The resulting strain, designated Vn14, exhibited no discernible phenotypic defects and replicated as rapidly as wild-type *V. natriegens* (Figure 1B-D).

### Induction of diverse large-scale genome rearrangements

To induce genome reconfigurations at the integrated *loxPsym* sites, we transformed the Vn14 strain with pMEV250, a plasmid encoding Cre recombinase under the control of an arabinose-inducible and glucose-repressed promoter (*26*). We adopted a population-based strategy to detect rearrangements (Figure 2A). A Vn14 population subjected to Cre induction was analyzed through comprehensive qPCR-based screening with primer combinations flanking *loxPsym* integrations to detect many possible inversions, duplications, and translocations (Figure 2B). This analysis revealed that a wide variety of predicted rearrangements was readily detected, with recombination events involving all chromosome segments (Figure 2C and S2).

**Figure 2.**
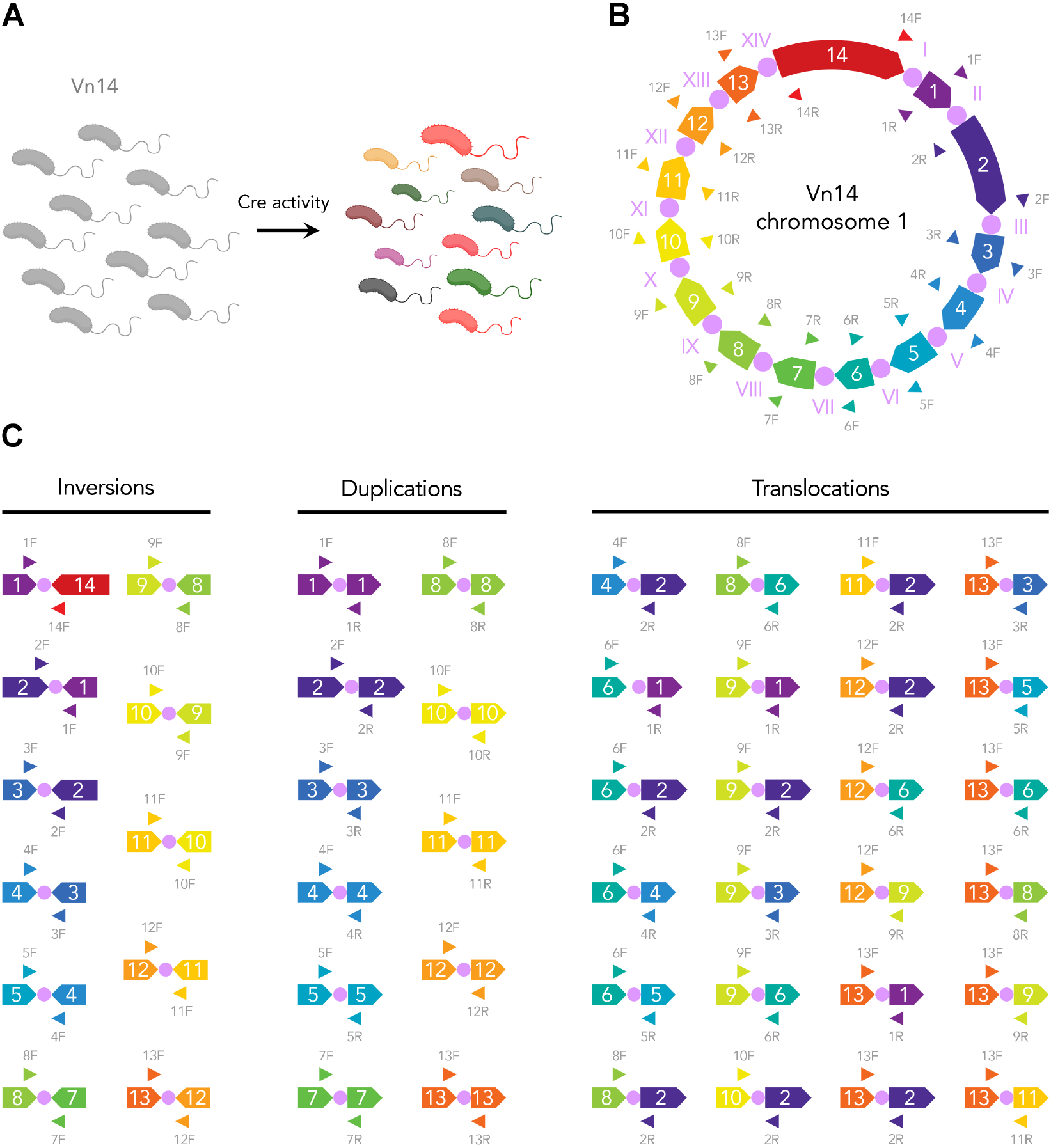
Induction of Cre activity in *V. natriegens* Vn14 generates a wide variety of large-scale genome rearrangements. (**A**) Induction of Cre recombinase activity in a population of Vn14 cells (gray) is expected to generate genetic diversity (colors) through large-scale rearrangements. (**B**) Chromosome 1 of strain Vn14 showing primers flanking *loxPsym* integrations (I to XIV) used for detecting rearrangements. Primers are represented as arrowheads. Each chromosome segment (1 to 14) is uniquely colored, and the direction of the pentagon represents the orientation. (**C**) A wide variety of selected possible rearrangements was detected using a qPCR screening based on combinations of the indicated primers.

### Evaluation of phenotype robustness to genome reconfigurations

To investigate phenotypic robustness to large-scale genome changes, we subjected a Vn14 population to induction of Cre activity under conditions to select for rearranged variants that retain rapid growth. We hypothesized that direct competition would remove rearrangements that lead to slow growth, allowing us to identify genome reconfiguration variants that support rapid cell division.

In addition to enabling MuGENT engineering, markers flanked by directional *loxP* sites that can be excised by Cre activity facilitate identification of SCRaMbLE events, as the loss of antibiotic resistance conferred by the marker would indicate cells that were effectively exposed to Cre activity. After 24 hours of growth, we screened for strains that had lost chloramphenicol resistance, presumably due to Cre-mediated excision of the *loxP*-flanked Cm^R^ marker, and selected these for further analysis. Screening of 4,397 isolated strains identified 167 that had lost resistance to chloramphenicol (Figure S3A). We then performed a qPCR-based screening using combinations of primers flanking *loxPsym* integrations to identify strains with potential rearrangements (Figure S3B). Further growth profiling of 18 selected strains (Vn14.1–Vn14.18) confirmed that they retained rapid growth, with strains Vn14.1 and Vn14.10 exhibiting ∼5% faster growth than the reference strain Vn14Δ*Cat*, lacking the Cm^R^ marker (Figure 3A). Whole-genome sequencing revealed a variety of large-scale genome changes that collectively spanned the entire chromosome. These comprised inversions of single segments (Vn14.1-9), two consecutive segments (Vn14.10-14), eight consecutive segments (Vn14.15), and one plus two consecutive segments (Vn14.16,17), as well as a translocation of a single inverted segment (Vn14.18) (Figure 3B). These rearrangements altered the GC skew of the replichores (the chromosome replication arms) (Figure S4), and several also displaced the positions of *oriC* and *ter*, thereby changing replichore lengths (Figure 3B).

**Figure 3.**
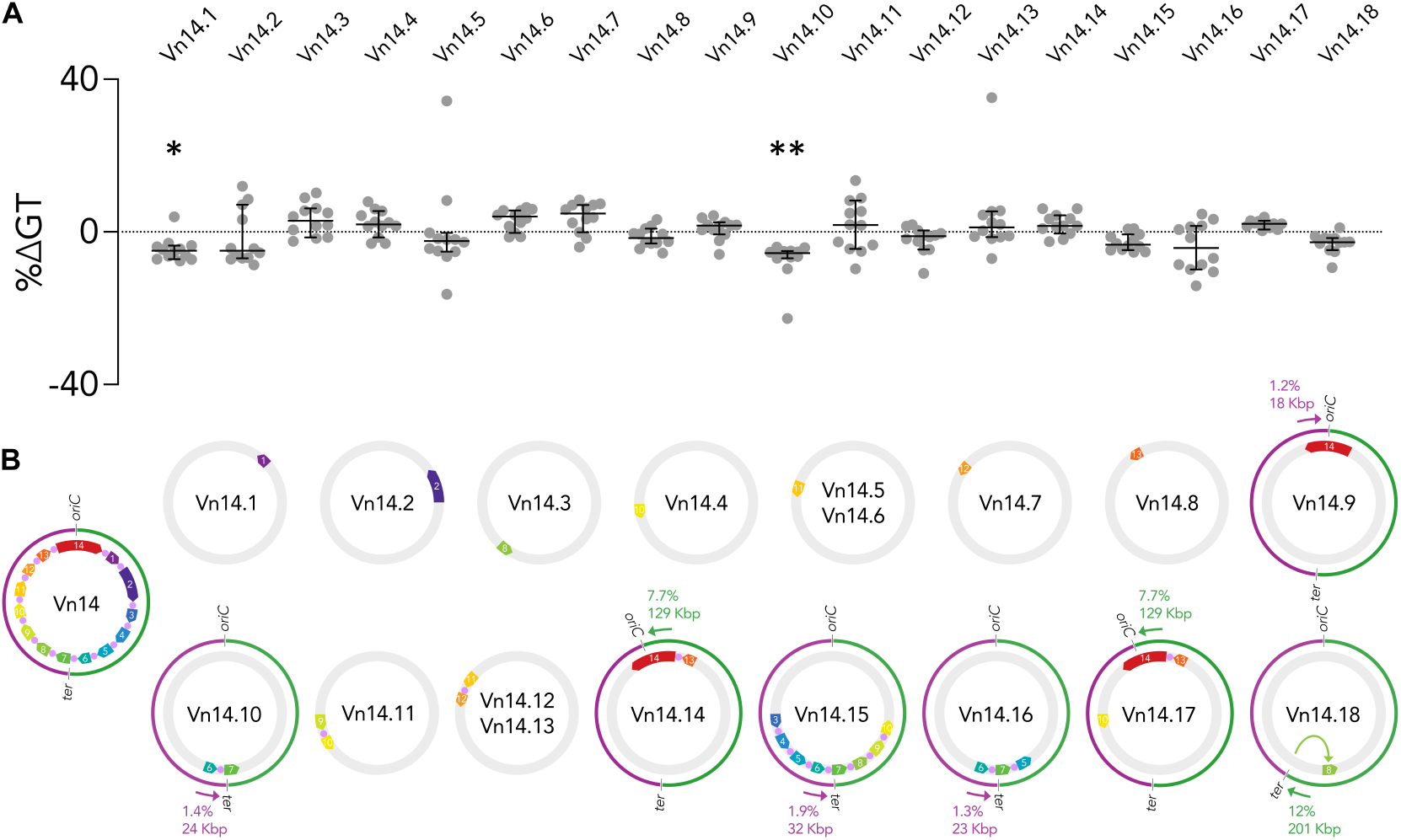
Evaluation of phenotype robustness to large genome changes. (**A**) Growth of isolated genome-rearranged strains that retain rapid cell division (strains Vn14.1 to Vn14.18), derived from strain Vn14 after induction of Cre recombinase activity. Generation time (GT) results are presented as percentage of variation (%ΔGT) of the median with 95% CI relative to reference strain Vn14 *Cat* lacking the Cm^R^ marker, as all rearranged strains analyzed (n = 12). (**B**) Large-scale genome changes in strains Vn14.1 to Vn14.18 shown in A. Each chromosome segment (1 to 14) is uniquely colored, and the direction of the pentagon represents the orientation. Chromosome replichores are shown as purple and green semicircles. Chromosome 1 of strain Vn14 serves as the reference, and genome changes identified in strains Vn14.1–Vn14.18 are displayed relative to it. Unchanged genomic regions are shown in gray. Arrows indicate how genome rearrangements have displaced the positions of *oriC* and *ter*. The percentage and Kbp changes in replichore length are also indicated.

## Discussion

We have established and evaluated the SCRaMbLE system to induce large-scale genome reconfigurations in bacteria. The design principles we defined for targeting *loxPsym* integrations in bacterial genomes resulted in a *V. natriegens* strain with no discernible alterations (Figure 1), suggesting that these principles can be applied to engineer other bacteria without causing significant defects.

Analysis of pooled Vn14 cells after Cre recombinase induction revealed an enormous diversity of genome rearrangements (Figure 2). This demonstrates the capacity of the SCRaMbLE system to generate and explore a massive space of genome reconfigurations in *V. natriegens* and offers new opportunities to study bacterial genome organization, function, and evolution.

Given recent evidence that natural selection broadly shapes gene arrangement along bacterial chromosomes, with the strongest positional biases observed in fast-growing species (*27*), we investigated how robust the growth of *V. natriegens* is to large-scale genome rearrangements. We found that many strains maintained rapid growth despite dramatic reconfigurations, indicating that distinct genome organizations can still support stable physiology, at least under our experimental conditions, with strains Vn14.1 and Vn14.10 displaying a gain-of-function as they grew faster than the reference strain Vn14Δ*Cat* (Figure 3). It is intriguing that such rearrangements are so well tolerated, given the tight linkage between chromosome organizational features and numerous cellular processes, as well as the strong conservation of bacterial genome structure along the chromosome *oriC*-*ter* axis. Bacterial chromosomes typically encode the majority of genes on the leading strand of replication so that their transcription is co-directional with replication fork progression, reducing the frequency of detrimental head-on collisions between replication and transcription machineries (*28-32*). We expected inversions would impact growth due to producing conflicts between the two machineries, which can be highly deleterious and result in poor growth (*13, 32-34*). However, isolated strains exhibited no substantial decrease in growth rates (Figure 3A).

Additional noteworthy features of these isolated strains are that their genome rearrangements also altered GC skew (Figure S4) and replichore lengths (Figure 3B). It would be interesting to investigate whether, and how, altered regions eventually acquire the characteristic GC skew of the rest of the replichore. Previous studies have shown that changes of less than ∼10% in replichore length have little effect on *Escherichia coli* growth (*13*). Consistent with this, the strains isolated here tolerated replichore length changes ranging from 1 to 12% without substantial growth defects (Figure 3).

Our findings provide a striking example of phenotypic robustness, suggesting that bacteria may tolerate chromosomal alterations more readily than previously appreciated. They further indicate that genome rearrangements may contribute to phenogenetic drift in bacteria (*35*), allowing the genotype to evolve through large-scale changes while the phenotype is preserved. Our work also demonstrates the feasibility of generating on-demand genomic diversity in *V. natriegens* through large-scale reconfigurations, opening paths to investigate the functional consequences of genomic rearrangements in bacteria, accelerate strain development for biotechnological applications, and illuminate evolutionary mechanisms that have remained underexplored due to the scarcity of genome-scale editing tools.

## Materials and Methods

### Strains, plasmids, media, and culturing conditions used in this study

Wild-type *V. natriegens* Rif^R^ (ATCC 14048) was used as the base strain in this study. *Escherichia coli* S17-1 was used to transfer plasmids to *V. natriegens* by conjugation. The plasmid pMMB-*tfoX*(Vc) was used to induce *V. natriegens* competence (*25*). The plasmid pMEV250, encoding the *Cre* recombinase gene under the control of an arabinose-inducible and glucose-repressed promoter, was used to induce genome rearrangements (*26*). Lysogeny broth (LB) was used as the standard medium for *E. coli*. LB3 (10 g/L of tryptone, 5 g/L of yeast extract and 30 g/L of NaCl) was used as the standard rich medium for *V. natriegens* (*18*), and LBv2 (10 g/L of tryptone, 5 g/L of yeast extract, 204 mM of NaCl, 4.2 mM of KCl and 23.14 mM of MgCl_2_x6H_2_O) was used to perform growth rate measurement assays. Antibiotic concentrations used for selection when required: carbenicillin 100-200 μg/mL, spectinomycin 100-200 μg/mL, chloramphenicol 5 μg/mL. Routine culturing of *V. natriegens* and *E. coli* was performed for 12-15 h at 30 °C or 37 °C with 200 rpm agitation in the case of liquid cultures.

### Desing and construction of V. natriegens strain Vn14

Genetic modifications were designed using Benchling software and the chromosome 1 sequence of *V. natriegens* strain ATCC 14048 (GenBank: CP101905.1). The Cm^R^ and Spec^R^ marker modules to enable MuGENT-based genome engineering were designed to integrate at the neutral *dns* locus (*25*). The Cm^R^ and Spec^R^ markers were flanked by directional *loxP* sites to allow Cre-mediated marker excision. Following the design principles described in the Results section, we designed 14 DNA constructs to target *loxPsym* integrations into the *V. natriegens* genome. All DNA constructs included ∼0.8-1.5 Kbp homology arms to allow targeted genome integration by homologous recombination. DNA constructs were synthesized by Genewiz and Gene Universal and are described in Table S1. The *loxPsym* sites were integrated into the specified positions of chromosome 1 through several rounds of MuGENT that swapped the Cm^R^ and Spec^R^ markers at the *dns* locus (*25*). The integration of *loxPsym* sites was confirmed by PCR genotyping using primers described in Table S2 and whole-genome sequencing. The MuGENT approach was also used to build the strain VnΔ*dns*Cm^R^, in which the Cm^R^ marker (*cat* gene) was integrated at the *dns* locus (see Figure 1A and Data File S1).

### Growth rate measurement and analysis

Growth was measured in LBv2 using clear, flat-bottom, 96-well Greiner plates with the lid by kinetic growth monitoring (Biotek Synergy H1 or TECAN Sunrise microplate reader) with continuous orbital shaking (807 cpm) for 18 h at 37 °C, and 600 nm optical density (OD_600_ nm) was measured every 2 min. Briefly, 12 fresh colonies per strain were inoculated in 1 mL of LBv2 in 96-deep well Axygen plates with a PIXL colony picker (Singer Instruments). Cultures were grown overnight for 15 h at 37ºC and 800 rpm in a 3 mm rotor incubator. Overnight cultures were then diluted 1/1,000 in 100 μL of final volume, and kinetic growth was monitored for up to 18 h. The slopes during the exponential phase and generation times (GT) were directly obtained using QurvE (*36*). The percentage of the GT variation of each strain was represented as the median +/-95% CI of the 12 independent colonies and was calculated for each colony as follows: % GT = [(GT-GTmedian^Ref^)/GTmedian^Ref^)] × 100. *Ref* corresponds to the reference strain used in each experiment, being the wild-type *V. natriegens* in Figure 1 and the Vn14Δ*Cat* strain, that is the Vn14 parental strain lacking the Cm^R^ marker, as all rearranged strains analyzed in Figure 3. All experiments were performed at least 3 times independently.

### Microscopy imaging and analysis

Cells grown overnight were diluted 1/1,000 and grown in LB3 at 37 °C with 200 rpm for 1.5 h and visualized using an Olympus BX63 microscope. Microscopy images were analyzed using ImageJ software (*37*). Only cultures that were grown on the same day for the same amount of time were directly compared and shown together.

### Growth on solid medium

Cells grown overnight were normalized to OD_600_ = 1 in LB3 and plated on LB3 agar following 10-fold serial dilutions, spotting 3 μL of each dilution. Plates were imaged after 24 h of growth at 37 °C using a PhenoBooth analyzer (Singer Instruments). Only cultures that were grown on the same plate for the same amount of time were directly compared and shown together. The last three spots of the dilution series are shown.

### Induction and detection of genome rearrangements

*V. natriegens* strain Vn14 was cured of plasmid pMMB-*tfoX*(Vc) by growth under no selection and then transformed with plasmid pMEV250 by conjugation from *E. coli* S17-1. Cells were grown overnight at 30 ºC and 200 rpm in 2 mL of LB3 supplemented with 100 μg/mL carbenicillin. Cultures were then normalized to OD_600_ = 0.1 and further diluted 1/1,000 in 5 mL of LB3 supplemented with 0.5 % w/v arabinose and 100 μg/mL carbenicillin. Cultures were grown for 24 h in 50 mL conical tubes at 30 °C and 200 rpm agitation. Genomic DNA was purified from 50-100 μL aliquots of overnight cultures by boiling samples for 10 min, cooling on ice for 5 min and spinning down at maximum speed for 5 min. Supernatants were analyzed to screen for genome rearrangements through qPCR-based screening using combinations of primers flanking *loxPsym* integrations (Table S2) that allow the detection of many possible inversions, duplications, and translocations (see Figure 2B and S2). The qPCR reactions were set up in 384-well plates using an Echo525 acoustic liquid-handling robot (Beckman Coulter) and carried out in a LightCycler 480 instrument (Roche) using iTaq Universal SYBR Green Supermix (Bio-Rad) as described in (*24*).

### Isolation of V. natriegens strains with reconfigured genomes

pMEV250-harboring Vn14 cells grown overnight as previously described were normalized to OD_600_ = 0.1 and further diluted 1/1,000 in 20 mL of LB3 supplemented with 0.5% arabinose w/v and 100 μg/mL carbenicillin. Cultures were grown for 24 h in 250 mL flasks at 37 °C with 200 rpm agitation. Grown cultures were serially diluted and plated on LB3 agar to isolate single colonies and incubated overnight at 37 ºC. Single colonies were replica plated on LB3 agar Nunc™ Omnitray™ plates with and without chloramphenicol selection (5 μg/mL) using a PIXL colony-picking robot (Singer Instruments). Loss of chloramphenicol resistance by Cre-mediated excision of the *loxP*-flanked Cm^R^ marker was used to identify cells effectively exposed to Cre activity (see Figure S3A). A subsequent boiled-colony qPCR-based analysis of these cells was used to detect potential genome rearrangements (see Figure S3B). The qPCR reactions were set up using an Echo525 acoustic liquid handling robot (Beckman Coulter) and carried out in a LightCycler 480 instrument (Roche) using iTaq Universal SYBR Green Supermix (Bio-Rad) as described in (*24*). Inversions, deletions, and translocations were screened using primers flanking all 14 *loxPsym* integrations, where only non-rearranged genome segments are expected to result in amplicon production. Duplications were screened using specific combinations of primers, where only duplications are expected to result in amplicon production. Genomic DNA of isolated strains was purified from cultures using the PureLink Genomic DNA kit (Thermo Scientific) and further analyzed by whole-genome sequencing (see Data File S2).

### Whole-genome sequencing and analysis

Genomic DNA was purified using PureLink Genomic DNA kit (Thermo Scientific) following the manufacturer’s instructions. Illumina sequencing was performed at Macrogen Inc. (South Korea). Long-read sequencing was performed either in-house using the Oxford Nanopore Technologies MinION platform or at Plasmidsaurus (USA). Genomic reads were mapped to the *V. natriegens* ATCC 14048 reference genome sequence (GenBank: CP101905.1) using Geneious Prime, the Mauve alignment algorithm and Benchling (see Data File S1 and S2).

### GC skew analysis

GC skew calculations and graphics were performed using Proksee (*38*).

## Acknowledgments

We thank Ankur B. Dalia for providing the V. natriegens Rif^R^ strain and plasmid pMMB-tfoX(Vc). MVB acknowledges support from a postdoctoral scholarship funded by the Ramón Areces Foundation. AGGO received a doctoral scholarship from Macquarie University and thanks the support of the Secretary of Internationalization and International Cooperation, UNSAM. LL is supported by a doctoral scholarship from CONICET. ITP acknowledges support for Macquarie University’s synthetic biology research from the Australian Research Council (ARC) Centre of Excellence in Synthetic Biology, the New South Wales (NSW) Office of the Chief Scientist and Engineer, the NSW Government Department of Primary Industries, and Bioplatforms Australia. ASB acknowledges support from the International Center for Genetic Engineering and Biotechnology (CRP/ARG18-06_EC to ASB), the Agencia Nacional de Promoción de la Investigación, el Desarrollo Tecnológico y la Innovación of Argentina (PICT-2017-0424, PICT-2018-0476, and PICT-2020-SERIEA-00521 to ASB), and the ECOS-SUD France–Argentina Program (A18ST06 to ASB). ASB is a career member of CONICET. BL gratefully acknowledges support from the Gordon and Betty Moore Foundation (GBMF9319; grant DOI: https://doi.org/10.37807/GBMF9319), the Ramón Areces Foundation, the ARC Centre of Excellence in Synthetic Biology, Macquarie University, Bioplatforms Australia, Twist Bioscience, the National Health and Medical Research Council (NHMRC), and the Allen Foundation.

