## Supplemental files for "Creating bacterial genomic diversity through large-scale reconfigurations reveals phenotype robustness to organizational genome change"

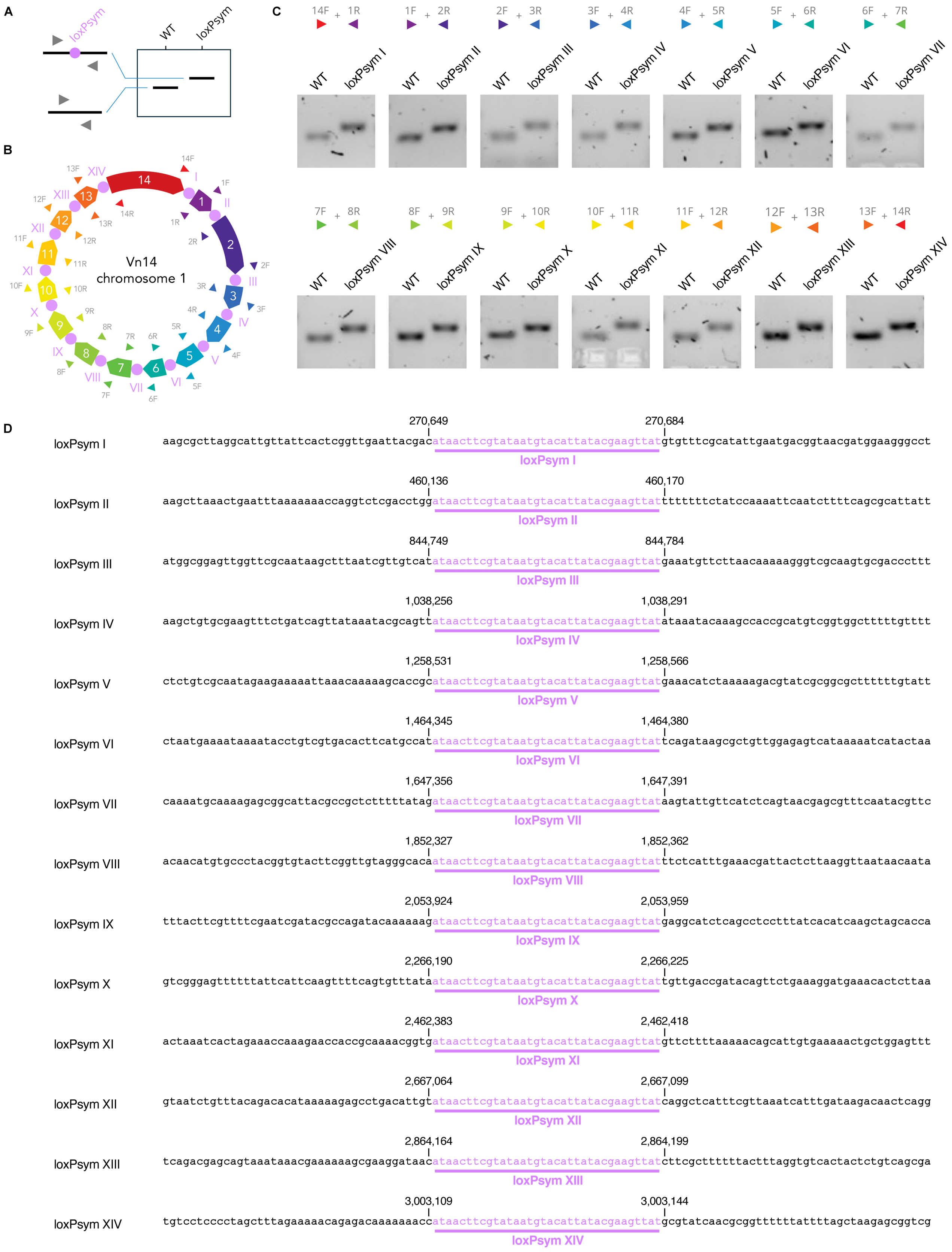
A
loxPsym
WT
loxPsym
B
Vn14
chromosome 1
C
WT
loxPsym I
loxPsym II
loxPsym III
loxPsym IV
loxPsym V
loxPsym VI
loxPsym VII
loxPsym VIII
loxPsym IX
loxPsym X
loxPsym XI
loxPsym XII
loxPsym XIII
loxPsym XIV
D
loxPsym I 270,649 270,684
aagcgcttaggcattgttattcactcggttgaattacgacataacttcgtataatgtacattatacgaagttatgtgtttcgcatattgaatgacggtaacgatggaagggcct
loxPsym II 460,136 460,170
aagcttaaactgaatttaaaaaaaccaggtctcgacctggataacttcgtataatgtacattatacgaagttatttttttctatccaaaattcaatcttttcagcgcattatt
loxPsym III 844,749 844,784
atggcggagttggttcgcaataagctttaatcgttgtcatataacttcgtataatgtacattatacgaagttatgaaatgttcttaacaaaaagggtcgcaagtgcgaccettt
loxPsym IV 1,038,256 1,038,291
aagctgtgcgaagtttctgatcagttataaatacgcagttataacttcgtataatgtacattatacgaagttatataaatacaaagccaccgcatgtcggtggctttttgtttt
loxPsym V 1,258,531 1,258,566
ctctgtcgcaatagaagaaaaattaaacaaaaagcaccgcataacttcgtataatgtacattatacgaagttatgaacatctaaaaagacgtatcgcggcgctttttgtatt
loxPsym VI 1,464,345 1,464,380
ctaatgaaaataaaatacctgtcgtgacacttcatgccatataacttcgtataatgtacattatacgaagttattcagataagcgctgttggagagtcataaaaatcatactaa
loxPsym VII 1,647,356 1,647,391
caaaatgcaaaagagcggcattacgccgctctttttatagataacttcgtataatgtacattatacgaagttataagtattgttcatctcagtaacgagcgtttcaatacgttc
loxPsym VIII 1,852,327 1,852,362
acaacatgtgccctacggtgtacttcggttgtagggcacaataacttcgtataatgtacattatacgaagttatttctcatttgaaacgattactcttaaggttaataacaata
loxPsym IX 2,053,924 2,053,959
tttacttcgttttcgaatcgatacgccagatacaaaaaagataacttcgtataatgtacattatacgaagttatgaggcatctcagcctcctttatcacatcaagctagcacca
loxPsym X 2,266,190 2,266,225
gtcgggagtttttttattcattcaagttttcagtgtttataataacttcgtataatgtacattatacgaagttattgttgaccgatacagttctgaaaggatgaaacactcttaa
loxPsym XI 2,462,383 2,462,418
actaaatcactagaaaccaaagaaccaccgcaaaacggtgataacttcgtataatgtacattatacgaagttatgttcttttaaaaacagcattgtgaaaaactgctggagttt
loxPsym XII 2,667,064 2,667,099
gtaatctgtttacagacacataaaaagagcctgacattgtataacttcgtataatgtacattatacgaagttatcaggctcatttcgttaaatcatttgataagacaactcagg
loxPsym XIII 2,864,164 2,864,199
tcagacgagcagtaaataaacgaaaaaagcgaaggataacataacttcgtataatgtacattatacgaagttatcttcgctttttttactttaggtgtcactactctgtcagcga
loxPsym XIV 3,003,109 3,003,144
tgtcctcccctagctttagaaaaacagagacaaaaaaaccataacttcgtataatgtacattatacgaagttatgcgtatcaacgcggtttttttattttagctaagagcggtcg

**Figure S1. Confirmation of *loxPsym* integrations into *Vibrio natriegens* chromosome 1.** (**A**) PCR genotyping strategy with primers flanking *loxPsym* integrations. Primers are represented as arrowheads. (**B**) Primers (arrowheads) flanking *loxPsym* integrations in chromosome 1. Each chromosome segment (1 to 14) is uniquely colored, and the direction of the pentagon represents the orientation. *LoxPsym* integrations are numbered from I to XIV. (**C**) Confirmation of *loxPsym* integrations as described in A and B. (**D**) Confirmation of *loxPsym* integrations by whole-genome sequencing.

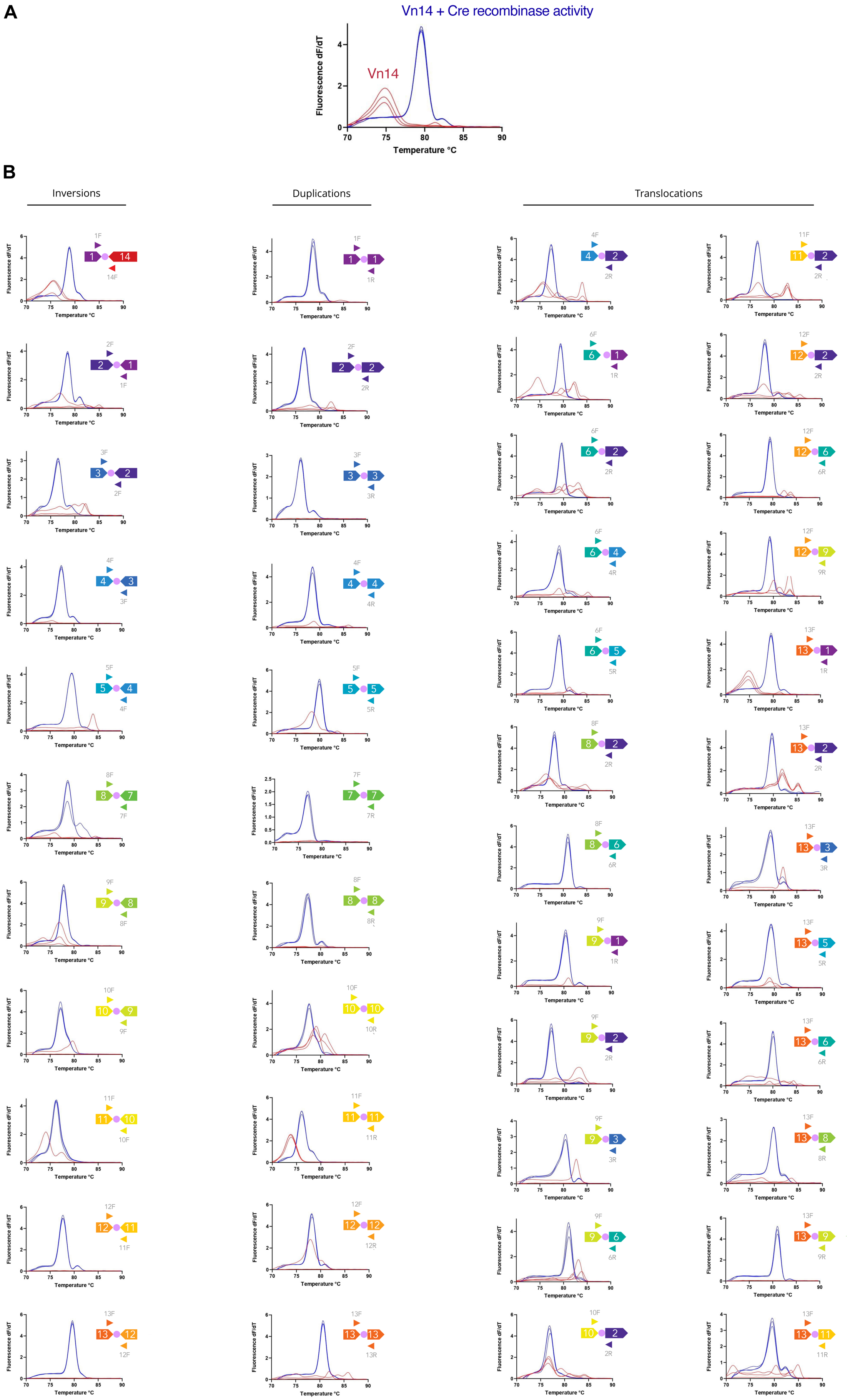
A
Vn14 + Cre recombinase activity
Vn14
Fluorescence dF/dT
Temperature °C
B
Inversions
Duplications
Translocations

**Figure S2. Detection of large-scale genome rearrangements in *V. natriegens* Vn14 after induction of Cre recombinase activity.** (**A**) Different combinations of primers flanking *loxPsym* integrations were used to identify genomic rearrangements through post-qPCR melt analysis. Only genome rearrangements at *loxPsym* sites are expected to result in amplicon production. A consistent peak in the derivative of the qPCR melt curve indicates amplicon production and is interpreted as detection of a large-scale rearrangement. A negative result is indicated by a flat signal (no amplification) or minor peaks derived from non-specific amplification. Data obtained from the analysis of Vn14 cells are shown in red. Data obtained from the analysis of Vn14 cells after induction of Cre recombinase activity are shown in blue. (**B**) Detection of selected rearrangements. Data n = 3. Primers are represented as arrowheads. Each chromosome segment is uniquely colored, and the direction of the pentagon represents its orientation. Pink circles indicate *loxPsym* sites. See Figure 2 for more details.

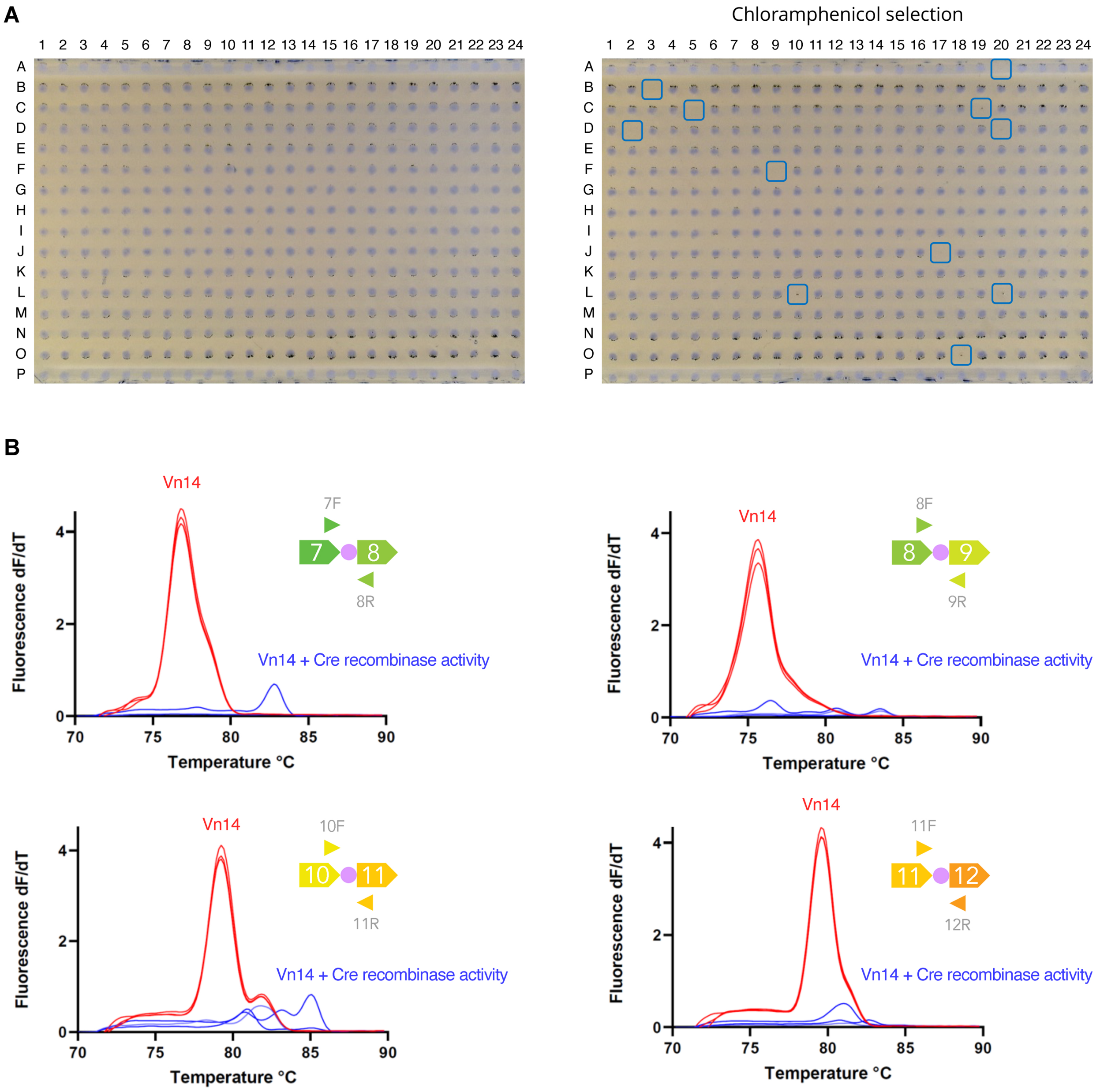


**Figure S3. Screening and isolation of *V. natriegens* Vn14 cells with genome reconfigurations.** (**A**) Representative photographs (inverted) of individual clones isolated from a population of Vn14 cells subjected to induction of Cre activity, replica plated with and without chloramphenicol selection. Loss of chloramphenicol resistance by Cre-mediated excision of the *loxP*-flanked $Cm^R$ marker enables the identification of cells effectively exposed to Cre activity, as indicated by the blue boxes. (**B**) A subsequent post-qPCR melt analysis was used to detect potential rearrangements. Inversions, deletions, and translocations were screened using primers flanking all 14 *loxPsym* integrations, where only non-rearranged genome segments are expected to result in amplicon production. Here, a consistent peak in the derivative of the qPCR melt curve indicates amplicon production and is interpreted as a negative result. A flat signal (no amplification) or minor peaks derived from non-specific amplification indicate a potential large-scale rearrangement. Duplications were screened using specific combinations of primers, where only duplications are expected to result in amplicon production (as in Figure S2). In this case, a consistent peak in the derivative of the qPCR melt curve is interpreted as detection of a potential duplication. A negative result is indicated by a flat signal (no amplification) or minor peaks derived from non-specific amplification. Representative data obtained from the analysis of Vn14 cells are shown in red. Representative data obtained from the analysis of isolated Vn14 strains after induction of Cre recombinase activity are shown in blue. Data shows potential rearrangements in Vn14 strains subjected to Cre recombinase activity. Data n = 3. Primers are represented as arrowheads. Each chromosome segment is uniquely colored, and the direction of the pentagon represents the orientation. Pink circles indicate *loxPsym* sites. See Figures 2, 3, and S2 for more details. Genome rearrangements were further confirmed by long-read whole-genome sequencing.

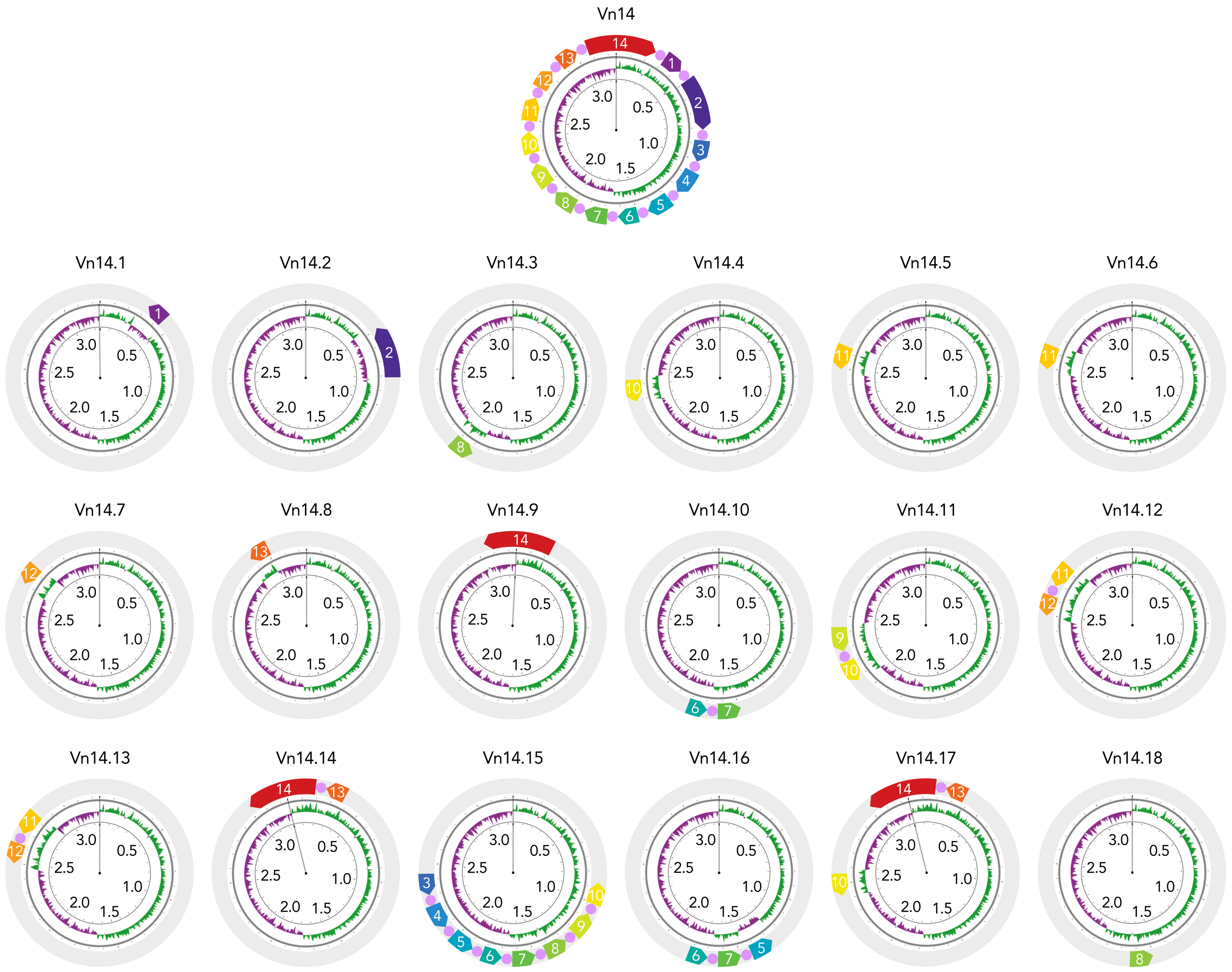


**Figure S4. GC skew alterations produced by genome rearrangements.** The outer circle depicts the structure of chromosome 1. Each of the 14 chromosome segments is uniquely color-coded, with pentagon orientation indicating segment direction. Strain Vn14 serves as the reference, and genome rearrangements identified in strains Vn14.1–Vn14.18 are shown relative to it; unchanged genomic regions are shown in gray. The central circle displays GC skew values along the replichores, with positive and negative skew shown in green and purple, respectively. The inner circle indicates nucleotide positions along the genome (Mbp). The dashed line marks the *oriC* position.

**Table S1. MuGENT constructs used in this study.**

Note *loxP* sequence is ataacttcgtatagcatacattatacgaagttat and *loxPsym* sequence is ataacttcgtataatgtacattatacgaagttat.

| Name | Sequence 5' - 3' |
|---|---|
| **SpecR marker** | Acgaacttgtaaagctgtaattgcggtaagagccgctgtttcggtgcgaagtacacgc<br>ggaccgagaagcgtctcttcaaattggtactcgcgcgtcatatcgatttcttcagctg<br>acaaaccgccttcagggccaatcagcaggcgcacttttttcaactggtgtaggcagggt<br>gttgatcgagtatttggcacgagggtgaaggttgagcttaagtccatcgtactcttct<br>ttgctccactcttccaagctcatgattgggcgaatttctggaacgatgttacgtccac<br>actgctcgcaagcactgatagcaatcttctgccattgggccagtttcttctcaaatcg<br>ttttgatcgagctttacgccacaacgttcagaaataaggggagtgatggtatttact<br>ccgagttcgactgacttctgaatggtgaactccatcttgtcgcctcgtgaaatcacct<br>gtcctaggtgaagatccaaaggggattcaatgctgttctctacgcgctcagagatgtc<br>tacgaggacattctttttgctgacttctgcgataactgcgggaaactctgcaccacta<br>ccgtcaaataggagaacttcctgaccttccttcatacgaagtacgcggccaatatggc<br>cagcggcgtcgtcacttaaagcgagtgtaccaagttggtgaatggtttctggatgata<br>aattcgagggatacgcatgagatttccagcaagagaataaacttagttcctaacatgg<br>atgctttccatcgaaaaaacaaggggcaagaggcgtttcaattggagtaaatgaaacg<br>cctcaaacagaaagattagcgattgtcgcgattggtgaggaataacttcgtatagcat<br>acattatacgaagttatatttaaatggcgcgccttatttgccgactaccttggtgatc<br>tcgcctttcacgtagtggacaaattcttccaactgatctgcgcgcgaggccaagcgat<br>cttcttcttgtccaagataagcctgtctagcttcacgtatgacgggctgatactgggc<br>cggcaggcgctccattgcccagtcggcagcgacctccttcggcgcgattttgccggtt<br>actgcgctgtaccaaatgcgggacaacgtaagcactacatttcgctcatcgccagccc<br>agtcgggcggcgagttccatagcgttaaggtttcatttagcgcctcaaatagatcctg<br>ttcaggaaccggatcaaagagttcctccgccgctggacctaccaaggcaacgctatgt<br>tctcttgcttttgtcagcaagatagccagatcaatgtcgatcgtggctggctcgaaga<br>tacctgcaagaatgtcattgcgctgccattctccaaattgcagttcgcgcttagctgg<br>ataacgccacggaatgatgtcgtcgtgcacaacaatggtgacttctacagcgcggaga<br>atctcgctctctccaggggaagccgaagtttccaaaaggtcgttgatcaaagctcgcc<br>gcgttgtttcatcaagccttacggtcaccgtaaccagcaaatcaatatcactgtgtgg<br>cttcaggccgccatccactgcggagccgtacaaatgtacggccagcaacgtcggttcg<br>agatggcgctcgatgacgccaactacctctgatagttgagtcgatacttcggcgatca<br>ccgcttccctcatgatgtttaactttgttttagggcgactgccctgctgcgtaacatc<br>gttgctgctccataacatcaaacatcgacccacggcgtaacgcgcttgctgcttggat<br>gcccgaggcatagactgtaccccaaaaaaacagtcataacaagccatgaaaaccgcca<br>ctgcgccgttaccaccgctgcgttcggtcaaggttctggaccagttgcgtgagcgcat<br>acgctacttgcattacagcttacgaaccgaacaggcttatgtccactgggttcgtgcc<br>ttcatccgtttccacggtgtgcgtcacccggcaaccttgggcagcagcgaagtcgagg<br>catttctgtcctggctggcgaacgagcgcaaggtttcggtctccacgcatcgtcaggc<br>attggcggccttgctgttcttctacgataacttcgtatagcatacattatacgaagtt<br>atagttaaagtctttaaaaagtatgactttatccattctatcactcagatattagctt<br>tcgtaataagcaagttaataataacaaaagaaatgttggtcaatatttcctattgaga<br>cttaattgaactgaggattaggaaagctggacaccagtgaagctaagaggctgacgac<br>atttttgacacgaatacacggcttgtttgcgcagcactttgttgtgacggcgaataga<br>aaggggatatgtcgtgcagcgacatcggtattcgaaggttttgccttgaacggaagag<br>acttcgaaactatgtgtggttttggccggaacgttaaatacggattccattacggttt<br>gccactctttgccatgaggacggacgcgaccgtaaacctgatatgtgattaagtgagc |

| | |
|---|---|
| | gacctcatgcggtatcacttcttgcaagaacgcttgggtgttttctttaaataaaacc<br>gggttaaggcgaatctcattgagctgaaggtacgcttttcctgctgctttacctcgga<br>ctttaaaggtcaggctgggaatagggaatgaacgcttaaaagcatgttcagctaatgc<br>gatatagtgcgccatcacctgttttgctttgtagcttagttcaatgtccattcgttcc<br>tctcaagttcaataaaaagcccccgcagttgcgagggccggatcatagcatcactgaa<br>ttaaggtgtgtgtttttttcttaatcgttgcatggtaggcgtgccaagtaccgtagccg<br>agaataggcatagtaaacagcataccgattccgtaggtggcgaaaccaattaagatac<br>ccccacagatgattgccgcccacacgatcattgctggaatgt |
| **CmR marker** | ccatgtagtggatttcgtagccgcggcgttgcgcttcaagcatcatggcaaagctaga<br>gtctttcttgatgttaatggacgaaattgggtccattacaataccgagtttgatcatt<br>atttttctccgttttaaccaagatcgccgaaacgaacttgtaaagctgtaattgcggt<br>aagagccgctgtttcggtgcgaagtacacgcggaccgagaagcgtctcttcaaattgg<br>tactcgcgcgtcatatcgatttcttcagctgacaaaccgccttcagggccaatcagca<br>ggcgcactttttcaactggtgtaggcagggtgttgatcgagtatttggcacgagggtg<br>aaggttgagcttaagtccatcgtactcttctttgctccactcttccaagctcatgatt<br>gggcgaatttctggaacgatgttacgtccacactgctcgcaagcactgatagcaatct<br>tctgccattgggccagtttcttctcaaatcgtttttgatcgagctttacgccacaacg<br>ttcagaaataagggggagtgatggtatttactccgagttcgactgacttctgaatggtg<br>aactccatcttgtcgcctcgtgaaatcacctgtcctaggtgaagatccaaaggggatt<br>caatgctgttctctacgcgctcagagatgtctacgaggacattcttttgctgacttc<br>tgcgataactgcgggaaactctgcaccactaccgtcaaataggagaacttcctgacct<br>tccttcatacgaagtacgcggccaatatggccagcggcgtcgtcacttaaagcgagtg<br>taccaagttggtgaatggtttctggatgataaattcgagggatacgcatgagatttcc<br>agcaagagaataaacttagttcctaacatggatgctttccatcgaaaaaacaagggc<br>aagaggcgtttcaattggagtaaatgaaacgcctcaaacagaaagattagcgattgtc<br>gcgattggtgaggaataacttcgtatagcatacattatacgaagttatatttaaatgg<br>cgcgccttacgccccgccctgccactcatcgcagtactgttgtaattcattaagcatt<br>ctgccgacatggaagccatcacaaacggcatgatgaacctgaatcgccagcggcatca<br>gcaccttgtcgccttgcgtataatatttgcccatggtgaaaacgggggcgaagaagtt<br>gtccatattggccacgtttaaatcaaaactggtgaaactcacccagggattggctgag<br>acgaaaaacatattctcaataaaccctttagggaaataggccaggttttcaccgtaac<br>acgccacatcttgcgaatatatgtgtagaaactgccggaaatcgtcgtggtattcact<br>ccagagcgatgaaaacgtttcagtttgctcatggaaaacggtgtaacaagggtgaaca<br>ctatcccatatcaccagctcaccgtctttcattgccatacgtaattccggatgagcat<br>tcatcaggcgggcaagaatgtgaataaaggccggataaaacttgtgcttatttttctt<br>tacggtctttaaaaaggccgtaatatccagctgaacggtctggttataggtacattga<br>gcaactgactgaaatgcctcaaaatgttctttacgatgccattgggatatatcaacgg<br>tggtatatccagtgatttttttctccattttagcttccttagctcctgaaaatctcga<br>caactcaaaaaatacgcccggtagtgatcttatttcattatggtgaaagttggaacct<br>cttacgtgccgatcaacgtctcattttcgccaaaagttggcccagggcttcccggtat<br>caacagggacaccaggatttatttattctgcgaagtgatcttccgtcacaggtaggcg<br>cgccataacttcgtatagcatacattatacgaagttatagttaaagtctttaaaaagt<br>atgactttatccattctatcactcagatattagctttcgtaataagcaagttaataat<br>aacaaaagaaatgttggtcaatatttcctattgagacttaattgaactgaggattagg<br>aaagctggacaccagtgaagctaagaggctgacgacattttttgacacgaatacacggc<br>ttgtttgcgcagcactttgttgtgacggcgaatagaaaggggatatgtcgtgcagcga<br>catcggtattcgaaggttttgccttgaacggaagagacttcgaaactatgtgtggttt<br>tggccggaacgttaaatacggattccattacggtttgccactctttgccatgaggacg<br>gacgcgaccgtaaacctgatatgtgattaagtgagcgacctcatgcggtatcacttct<br>tgcaagaacgcttgggtgttttctttaaataaaaccgggttaaggcgaatctcattga<br>gctgaaggtacgcttttcctgctgctttacctcggactttaaaggtcaggctgggaat |

| | |
|---|---|
| | agggaatgaacgcttaaaagcatgttcagctaatgcgatatagtgcgccatcacctgt<br>tttgctttgtagcttagttcaatgtccattcgttcctctcaagttcaataaaagccc<br>ccgcagttgcgagggccggatcatagcatcactgaattaaggtgtgtgtttttctta<br>atcgttgcatggtaggcgtgccaagtaccgtagccgagaataggcatagtaaacagca<br>taccgattccgtaggtggcgaaaccaattaagataccccacagatgattgccgccca<br>cacgatcattgctggaatgttggatttgaccgcattaaagctggtgaataccgccgtc<br>atcatgtcaactctgcgctccatcattaatggaatagagaaggcagaaatgctgaata<br>ccacgcttgctagtacaaaaccaatgagtgacccaatcactaagaagggcaa |
| ***loxPsym* I** | actccctgctctcgcaaatgtccgcttccaacctgctgggcatatattgggctctggg<br>tatgtcgaatttaaactgtcaaataatgaagttgtcgttttctccggtgacttagggc<br>catcgaatacgccacttttgcctgatccacaatcgccagagcgcgcggactacctatt<br>tatagaatccacctacggcgatagacaacatgaagacattgcaacacgtacaaagcgt<br>ttaaacgacatcattactcatgcgttgaaagatggtggtgattctgatccctgctt<br>ttagtgtcggtcgaacacaagagctgctatttgatatagagcaattgattaaccagca<br>tgaattggcatcgtcattacccgtcatcctagactcccctctagcaagagaagtgaca<br>aaaacgtatcgacgttttaaaccttatggagtcaggaagcaaagcaaagattagaca<br>atcaacgccatccacttgctttcgaccagtgtattacagttgaaagtcacaccgagca<br>taaagccctagtaaaccgactcgcttcaactgatgaacccgccattgttgtagccgct<br>tccggaatgtgtgaaggcggcagaatcgtcaattatttaaaagcacttcttcccgatg<br>aaagaaacgatgtgctattcgcaggttatcaagctcaaggtacgttaggtgaggcgat<br>acaatcaggcaattcagaggttgagatagatgctcaagaaatccaggttaatgctcag<br>attcatactatttctggttattccgcccatgcagacaaaagtgatttactcaagttta<br>ttactggcattcccacacaaccaaaagctattcatttgattcatggggaagacgaggc<br>ccaacaagcattatgcaaggagttagagaagaaaggatttaaggtggtttattgatgg<br>cggagtttataagacctaagacttaagtacctaggcccttccatcgttaccgtcattc<br>aatatgcgaaacacataacttcgtataatgtacattatacgaagttatgtcgtaattc<br>aaccgagtgaataacaatgcctaagcgcttacaaggctatccaccttcgttaagtttt<br>ccaattcggtttggttaacctctgtttggcttacctggctctggttccaaactaaatc<br>actaatcccatcggtagggttgaagtcagttggagcggccaataggagctctcgaata<br>cggacgtgatccatgttatgacaagcaaaatccagttcatctagtaaggattcatatt<br>catcaagaggtaagtaaacttcatgggcagtcatgatccgctcatgggaagtgttctt<br>aacgttatcgccaatcagtaattcctcgtatagtttctcaccaggtcttaggcccgca<br>aacagaatttcaatatcaccatgaggattcgattctgatttcacttcaagaccagaaa<br>ggtggataaggttgctcgcaagatcggtaattttaactggttcacccatgtctagtac<br>gaaaacatccccgcccttacccatggctccagcttgtatcactaattgagccgcctct<br>ggaatggtcatgaaaaagcgcgtgatttctgggtgtgtcacggtaactggaccgccac<br>gttcaatttgctttttaaacaggggaataactgaaccagatgagcctaatacgttgcc<br>aaatcgaaccatacagaaacgtgtaccattaggcttttattttcacgctccgcgagt<br>gcctgtagcccaagctcagccatgcgtttggttgtccccatgacattggttggacgca<br>cagctttatcagtcgaaatcagaacaaaagactcgactcctgctttgatcgctgcgca<br>tgccgtgtaataggtgccatagatattatttcgtacgccttctacgacgttgtattcc<br>actaaaggaacatgcttatacgcagcggcatgataaacggtttccaccttgaacgatt<br>tcatcgccgtttcgaggcggtgctggcgctggacagaacctaatagaggaatgatatc<br>gcaa |
| ***loxPsym* II** | gcttcaaatgtcggctcatagttattaatttcttttacaatttttctaaggcggcgca<br>gtgttcgatcgttgcgactgccaataacttttgtcagtagcttagttatcatttttcg<br>tgaatctctctttcatcagaccgttcccgatctactcgtaatgccttcagtctctgcc<br>agtttggtactaagaatactgatataaatcatttaacaaatgatattgagtccaaaca<br>tccatatttcaaggctaaatttagaaacattggggcttagtgtaaggattttaatcg<br>atacaaccataaatagacctatttgtttgaagcactctgtaaattttattgaaagtgg<br>tgatagattagccaaatttttatttgtaagtctccaaatctatcaatttgctttcatt |

| | |
|---|---|
| | atgcggcgtccaaatcaattgtaataaagacgctaatgtaaactatacctagatagaa<br>agctgctctagaatagcggtatatccacaaaattagattttacctatgcgtgatcatc<br>gaccaacctcaacagaggatgttattgctgaatctcagtttaagaagattcaagagca<br>tgctggagaaatcttgcagctcaacaaagccttacaggatatcttgcctaaaggcacg<br>gctgatcattgccgagttgctaatgtccgtaacggtcatctattgatcgatgtgtcta<br>gcgctgccatcaagatgaagatcgactacgatcgcctaatgatcctcaataaactacg<br>aactcaaggctatgcaaagttgatcagcgtggatgtccgaattaatccatccctttat<br>cgcaaccggtacgagcaagacgatagacccaagcgaccgccgttaactgagtcagcgg<br>ccaaatcattaatgattattgcggatatggcgccacctaaaattcaagagcgcctgaa<br>aaggctggctgacatggctgataagtcaaagtaataatgcgctgaaaagattgaattt<br>tggatagaaaaaaataacttcgtataatgtacattatacgaagttatccaggtcgag<br>acctggtttttttaaattcagtttaagcttatgctaataccatatttggctctgcaaa<br>tgcgactggagaacccgcttcttcttcgaatgtagtccattcccatgcttcctgatca<br>gccagaacagcacgtagaagttggttgtttagaccgtgacctgatttgaatgcacgga<br>actcaccaataattgggtgtccacacatgtaaaggtcaccaattgcgtccaatacttt<br>atgagtaacgaactcattatcaaaacgcagaccttcttcattaagaattcggtaatca<br>tccagtacgatagcgttatcaaagctaccacccaaaactaagttctgagactgtagat<br>attcaatatcacgcataaaaccaaacgtacgtgcacgagaaatctcgcgaacaaaccc<br>tttagatgagaagtcaaacaacaggcgctgctcgtcagactcaattgcagggtggtta<br>aattcaatctcgaagtccatacggaagccgttaaatggaacaaactctgcccacttat<br>cgccatcttcgaaacgaactggtttttaatgcggataaaacgcttgggagcattttg<br>catttcgatacctgcttgttgaagcaagtatacaaatgggcttgcgctaccatccata<br>attggaatttcaggagcgtctacttcaacaacaatgttgtcgatacccataccagcga<br>gcgcagcattcaagtgctcaactgttgaaatacgcacgccttcgtcgttaactaatgc<br>agtacatagcatagtgtcacgcacagacgcaggatcagctggaaaatctacaggtgga<br>tttacatctgtacgacggtaaatgatacctgtatttgcagcagccgggcgcagagtaa<br>gcgtgactttacgaccagagtggagacccacaccagttgttttcactatttctttcag<br>agtacgttgtctgatcatctgcttgcctctctaaagtgtgctacaaaacacggtcgtc<br>tgta |
| ***loxPsym* III** | tgatggcgcatttaccatcttctacatgggtatcaaccttggcgctctgctcgctggt<br>gtcgtttctggctcggtgaccaactcattcggctggaaagctggtttcgttgctgcgg<br>gcatcggtatgcttatgagccttgtgatgcaaataagcttcgctcaatcctggctagg<br>tgatatcggccgcgaaccagcagccaaacgcgcactagcaaatcagaaatcaaccaag<br>aagcagccacttactaaagaagaagtggaccgcattaaagtcattctggtcatgagtc<br>ttttcactatcgtattctgggctggctttgagcaagcaggtggcctgatgaatatcta<br>cacccagcaatacactgaccgtatgattggcagtttcgaagtccctgcggcttggttc<br>cagtcacttaacccgttctttattatcacgcttgcacctgtgcttgcggtgctgtggg<br>ttaagctaggcaaacgtgagccaaactcaccagtgaagtttgcaatggcgatgttctt<br>cctggctctgggtttcctatgtatggttggtgcggtacttgagcaaggcggtgatacc<br>acagtgaaaacatcaatgctgtggttagtgggtgctttcttcttccatacactgggtg<br>aactttgcctgtctccaatcggcttgtctctggtaaccaaacttgcgcctctgcgttt<br>ggcatcattaatgatgggtgcatggttcggctgtaatgcgattgcaaactacgtagct<br>ggctatgtgggatcccatgttggtgagctaggtgcgatggctatctttagcggtatcg<br>cagcgagcgcgacggtcagtggtctaattttgctgttgttctctaacacgctcgtgcg<br>ctggatgcacggtgctgaaagttcagcgagcaacgcggaacaagtagaagagcaacaa<br>gtacaagttgcttaatacatcattcaaactaaaaagggtcgcacttgcgaccctttttt<br>gttaagaacatttcataacttcgtataatgtacattatacgaagttatatgacaacga<br>ttaaagcttattgcgaaccaactccgccatcatctcaatatgtcgctcatcatcattc<br>aggcaaggaatgtaattaaactgctcaccgcctgcttctaagaaaatgtgtttgttct<br>ctcccgcgatttcttctagcgtttccaaacaatccgaagaaaatgctggtgtcatgat<br>atcaatctttttaatacctttggcaggcaacgtttccagagttttatcagtgtatggc |

| | |
|---|---|
| | tgtaaccactcttcacgcccaaaacgggactgataagccatccctatctgcgtttcat<br>tcaacccaagtgcttcacccagcaagcgagtcgtcgtttcacaatgttgaggataaat<br>gtcaccgttatctgcaaaacgtttcggtatgccgtgataggaacatagcagatactcc<br>ccttgtccatgttcagcccaatacgctcttacggattgagccaaggcatcaatgaaca<br>ttggatggtcgtgataatcacgaattaagctatatgccggtatgacgggtaatttctt<br>aaaagctttggttatcccatcagacaccgccgctgttgttgtacctgaatactgagga<br>tacaaaggcaatacgatgacctcatcgaccccttgcgatagtaaggcttcaaacccg<br>cgtgtaaacttggattaccataagtcatgccaagctctacgggcatatctaactgctt<br>agccagtttcttcgcttggcgctttgagtacaccatcagaggggaaccctcttccatc<br>caaacggactcataaagcttggcaacttttggcgcacgtattggcaaataaccccgt<br>gcaaaattggacaccaaagccatcgtgtcatgtcgaccactcggtgatcatgaagaaa<br>ctggctcaagaagcgctttacagcaggagcagttggttcatcaggtgtacccaagtta<br>accagaagcacaccttgtttttttatttttgcatagcgtccctttgtttccagcgtg<br>tacc |
| ***loxPsym* IV** | gatggcaaaattcaggtctacgttcaacactacttgtttgcagagacaaccatggatt<br>tgcgtgcgccgggtgtcagagaaattcgttttgagcctaaacccgttgaacaacttgc<br>aaacgaaatgggtgaagctgacggaaatgttcaggttggtaacttagccgatgtatcg<br>ccaaccgtgactgaagagaagtttcttaagacttaccgccttgatcaaaaacgccgca<br>tgcgaagcagcgaaacacattacctggacagccctttaatggggatgatcatccaggt<br>acgccgtgtaaactaaaccataaaaataataaaagcgcagctcaggctgcgcttttgt<br>tttataactaaaaaacgaccagtaagggagtcatatcaattcatgagccccgatttttg<br>aaattattacgccgcgcttagctttacgtcttattcctgccgatgatgcaaatgctct<br>gcatacgctgcttgccgaatctccttcactgcatacctggttagactggtgtggtgaa<br>aacgtgtcactcaaatacgcacaggacttcttgctttccacacgcttgaactgggtga<br>aaacagaagcttttggttttggtgtttatgaccgcgaaagcaatgcccttcttggcat<br>ggtggctattaatgagctctaccatacctttaacatggcgagtattggctattggatt<br>gccgaccgttatcagcgacagggttacgcacaagaagcggtttgtgcattagccgaat<br>tctgtttcgctaaacttagcctgacgcgtatcgaaattgtctgcgatccagataatac<br>tgccagtcaggcactgatcgaatccgttggagcacaccaagaggcaattgccagaaac<br>cgctacatctttgacggtaagccaaaagatggcattgtctattcattgctaccagaag<br>atctaaagtaacgtttgggacgcaactgcagaaaaacaaaaagccaccgacatgcggt<br>ggctttgtatttatataacttcgtataatgtacattatacgaagttataactgcgtat<br>ttataactgatcagaaacttcgcacagctttcgctaagcgaagtttactctattaatg<br>cagcttcaaattaggacgcaatacgcggttgattctaccaaccagcatcatcaaacca<br>gtcttgaaaaggccgtgcaacgcaatttggtgcatgcggtaaagtgaaatatacacca<br>cgcgggcgatacgaccctcaaccatcattgagcctttggttaagttgcccatcaagct<br>gcccactgtcgagaactggctcagagaaaccaaagaaccatgatctttgtacacatag<br>tcttttagttcacgaccgttgatcatcgcgacaatgttgctgaatgctcggcttgcca<br>tttggtgtgcagcttgagctcttggcggtacaaacgaaccatcttcctgagtacactg<br>agccaaatcaccgatcacaaagatgttgtcatcacgcgtagtttgaagggtactttc<br>accacgagctggttgattcggttcgtctccagacccgcgatgtctttgatgaagtcag<br>gcgctttaatacctgccgcccaaaccatgattttagcggaaatcttctcaccgtcttt<br>ggtcgttaagccgtctttctcagcttttgtcaccatcgttgcggtacgcacattcacg<br>ccaagtttcgtcaattcttgatgtgctgcagtagagattcgagaaggaagcgcaggga<br>gaatacgtgggccagcttcaatcaagtttacattgagcttgctcgagtctaagtcacc<br>aaagccgtatgcacgaagctctttaatcgcattatgcagctcagctgaaagttcaaca<br>cctgttgcacctgcaccaacgattgcgatatcaacggtaccgttaccattttcgcgt<br>ggagcttaagaaactcattgttcatttcgctgcggaatcgatgagcctgttcagggct<br>gtcgaggaaaatacagttttcacgaacacccggagtattgaagtcgtttgaagtagaa<br>ccta |

| | |
|---|---|
| | aatcacttctgtttcaatttaaacccggtcagtttatcaacctgggtgtcgaaattga<br>tggaaaattggagtttcgtgcctattcaatcagctctgttaatgaagacgatcacctt<br>caactcaccatcaaacgtgtagctagcggtaaagtatcgaactacatcgttgactcgc<br>tgctgctcggtgataccgttcaagcattaccaccagcgggcgaatttaactgcattga<br>ccacccacccgttttacgtgatggcgaaacaaaagccttgcttatcagtgctggttgc<br>ggcgtcacgccagtattttcgatggcaaaacactggttgagtaatcagggtgagaatg<br>acatcgatatggatattgaatttcttcatatcgcgcgtagcccagaagagacgattta<br>ctacgatcagctagaaacgttcggtgccgtgtatcccaatttccatttgaagctgttg<br>ctgaagaacagcgaaggaacgttgcatccgcaaggtcgcttgaacgccgactggctgc<br>aaacgctcgtccctgattttaagcaacgcacggtttacctctgtggcccaagccagtt<br>catgcaagatgttcacggatacttgagcgaactcggttttgacatggcaaacttctat<br>caagagagctttacgccagcagctacggaaccttcagcggtatcagatgaaaccgcca<br>gtgtctccgtgcctgactttgctcaaaccattgatgctcaaaaaggtcaggtattggc<br>agatgttcttgaaggtgcgggcttacctctcatcatcgcatgtcgaagtggcatatgt<br>gggtcttgcaagtgcaaagttcgtaaaggcagcgtctcttccaccagcctagaaacat<br>tgacgcaagaagaaattgaacaaggctatgtactcgcctgctcatcaaccattgaagc<br>agatcttgaagtgcagattggttaattttccaaatacaaaaaagcgccgcgatacgtc<br>tttttagatgtttcataacttcgtataatgtacattatacgaagttatgcggtgcttt<br>ttgtttaatttttcttctattgcgacagagagtttttagctgaatagcaacactttc<br>aatgccgcttcgctgtccacgtccgtaaactctggcgggttatccagacgctcgcgga<br>acgtcagttgtggcgcttcttcttgcatgctttcaatcaagaaatcactggtcactga<br>tggtgagttcacacacgccatcacttgaccgccttctgccaacaaatcaggcaatcga<br>cgcagaatctttttgtagtctttggtcagtgcaaagctgcctttctggaacgaaggcg<br>ggtcaatgatgatcatatcgtacgggccagatttcttaattttgccccacgacttaaa<br>gatgtcatgccccataaagctgacttggtttgtgttgtgaccatttagcttatggttc<br>tcgcgccctttagacagcgacgctttcgccatgtctacgtttacgactttatctgcgc<br>cacccgcaatggcagcgacagaaaagccacaggtataagcaaacaagttaagcacgtt<br>tttatgcttggcatttcgcgtacccaatcgcgaccgtaacgcatgtccaaaaacaga<br>ccaaagttttggttgcggccaatatccaactggtatttcaaaccgctttcttcaacga<br>cggggcgagcatctaagtccccaacaagcacttctgaaggtgcgccatcggcataacg<br>gtgctgaagtaccattgtacggccttgtttacttgcccacagatctgattcagctaac<br>gcattgaggccacttttcaattcagcgagaaacgcgtcgtcgacttctttgaataagt<br>taatgatcaactgaccttgtaaccagtcgcaagtgatttgctctagaccttcaaaacg<br>acggccacggccgtgaaacagtcgacgaatttcattcggagcttgcgcgagctcggtt<br>tccagatgctgaaaaaacagcgaaagttgtgatgcttgcatcgaattctcttattaat<br>ttct |
| ***loxPsym* V** | (continued above) |
| ***loxPsym* VI** | tgactcgtttcgttaccatctgacgccaattacaatagaccacaaagttatcttcaca<br>gttaccatgagaagaagtgatcagctagtggagccttttgtaaactagctaattgtgt<br>tattcatcattgggaagaggaatatccatatttagtcgtaaaacaatagagatctaat<br>aaaagaaggatagaataaatttttaaaaagtgtggcagttagtccagatttagtattc<br>gctggtgagtgacaggttaatcgataagttagacctgtttcgctctttaggatctaga<br>ccccgtatcataggacaactctcagatatgagagcttagtttgggtaagaaacggaag<br>ttacttacgtgtttaatatgaaacttcactcctatacttcaacatcttatgcattaat<br>gccctcggttcaaaaatatattcctgaacacaccactcctttcatcgagcaatatgct<br>gctttggactttaatttacttggaagccaaatttgttgtagagacgaatctgcatgta<br>attttattgtcgatggaaacgagccaagtatgggaaaaatagtcaagtttacacgaaa<br>caacttgaattcgatcagctttgcttttgattctttcggcgaactcgtaatcaccaaa<br>tggtcacacaaccgaagggatctgaagttcttctggcatctcgtttcccatatctgag<br>tgataaccagcggcatgcggtactggcgacaacagaaatcgagtctaactaccctatt<br>attaatgattctcgtggctagggtcgtctgaacttggttgacgcagcggatagtttcg<br>gtgcttttaatagcaatatttatgtatatatggatgctgatgaaggtggatttaacac |

| | |
|---|---|
| | agaagactactggcgtaatgatatctctggtgaaggtcgcctggagaaaagtggtaca<br>aatggcagcgtatcaatgaagtcgctgccatttagtatgattttatgactctccaa<br>cagcgcttatctgaataacttcgtataatgtacattatacgaagttatatggcatgaa<br>gtgtcacgacaggtatttattttcattagcttactgattcggcttgctgttggcgca<br>tctgcagtcgactcaccaaccacatcacaaagccaccagcaaataacatggctgccac<br>ataaccggtgtaattggcactataaccattgataaacactaagctacccgccaatgga<br>ccaatcgcattagccgagttaaaggcacactgcagtaatgccccaatcaccgcgtgtc<br>cccctttggcaacacgcatcaacatcgcttgtatcactgtgcttaagcctaggctcgc<br>acccaacaaaaataccaccagatacaagaaccaaagattatcggcagccgatacgtac<br>agaatggctagcaacacacaataaactaaagccacgcctgtcactttcgcgactaccc<br>tatcgtgcacatgaccaaacaaccagttgcccaaagtcgatcccacaccaaaaatcac<br>catcgcaatcgagatgtagtattcaggcacttgagttacatcaatcatggtatcagct<br>agataagtatacaggcagaacacgccaccaaagccgatcaccgcgatgaatacaattg<br>gccagatcagcggattttcagtactgccaagtcacttttagcccctcgccttcagc<br>ctgcagcgaaggcaatacacgccacaacagcacacaagcgacgaaagctccaattgaa<br>accagcaacaacgcccaacgccagctaatgttttgcgccaataatgtcgctaatggca<br>caccgacgatagtggcgatggtaagaccagaaaatacccgcgacatatataacgtact<br>tcgtccctctggagccaggcgagcgccaaaaagcaacgccgtaccgaaataagcacca<br>tgaggtaagccacttaaaaagcgaaacaatacgatttgctcaaacgaggttgcggcag<br>cgcttaaaccattaaaagcaaaaatcaaaaaagcgaacaacaccaaagcactgcggcg<br>attc |
| ***loxPsym*** **VII** | tggctctgcttacatggctcgtgaggaaggcgcgaacattgatatcatctggccagaa<br>aaaggcacaatcttctggatggacagtcttgcgattcctgcgggtgctaaaaacgtgg<br>aagccgcacataagatgatcgacttcctgcttcgtccagagaatgcggctaagattgc<br>actggagattggttaccctactccagttaagacggcgtttgatctacttccaaaagaa<br>tttgttaacgaccaaagcatcttccctccgcaagaagtgatggacagcggtacttggc<br>aagatgaagttggtgaagcgagtgtgatgtacgatgagtacttccaaaagctaaaagt<br>aaacaactaataccagtcaaagtataaagaaaaaagcggccaactaaattggccgctt<br>ttattgtttttgaaggctttatttgtcttccaaagataaaatttcattcaccaacttt<br>ggtacttcgatacttgccttgccgtagcgtttctcttcgaactcactttctattgcac<br>ttggctccagattaatctcgatggtatgtgctccgtgcatttttgcatcgtggacaaa<br>cccagccgcaggatacaccacaccagatgtaccaatagaaataaacaagtccgcttgt<br>tctagcgcttcgtaaatatcacccatgcgcagaggcatttcgccaaaccaaacaatgt<br>gagggcgcatttgggacggaatttgacaacagtggcaaagctcacccgtttcaatatc<br>ctcagtgtgttctatcacctgatgcgattcgctacagcgggctttgagcagttcgcca<br>tgcatgtggatgacattgtggcttccgcctctttcatgcaaattatcgatattttgcg<br>taatgacggtcactttgccatccagctctgcctcaagtcgccctaaagcaagatgcgc<br>ggcgtttggctgtatcgcctcgtcctgtagcttttacgacgttgattataaaatgcc<br>tgaacaaggtcgggatctcgctcgaatccttcaggggttgccacgtcttcaatacggt<br>ggttctcccaaaggccatcttgagctcgaaatgtttgaatgccggactctgctgagat<br>acctgccccagtaagtactacaatgtttctatacgggaaattcataactgatccttgt<br>tgttctttttcattagcttaacactgaaacaatatcagctgtagtaatatggattaga<br>caatggtagtaatgaacaataataaaatgatattttgagaggaaaagcaatgagacac<br>gtatctctggctcttgtcttacttacgactctagcgggctgtcaattgaccaatgttg<br>aaggtgaaattgatgatgtgagagtgagcgttggtaccaatgatcatgacagacacga<br>tggtcatggtaaattctgccctccgggtcaggcaaaaaaaggaaattgctagaacgta<br>ttgaaacgctcgttactgagatgaacaatacttataacttcgtataatgtacattata<br>cgaagttatctataaaaagagcggcgtaatgccgctcttttgcatttgttaatcttc<br>gttagttgaagagatttgtggatagctaagtcggcaccattaaattcatcttcttca<br>gacaatctaatgcccgtaactgcgttaattgcgccatagacagccaacgcacctaata<br>cagcaatagaaatacctaatacggttcccataagctgaacaacaaggctaacaccgcc |

| | |
|---|---|
| | taacccgcctaacgctgtttgtccaaagatgcccgcagcaatgcctccccatgcgcca<br>caaataccgtgaagaggccaaacacccaacacatcgtcaatcttggttttgttttgta<br>gatgtgtaaacaggtaaacaaagagcacgcccgcaatcgcacctgtcaccagcgcacc<br>gattgggtgcataaggtcagagcccgcacacacggcaactaatccagccagagggccg<br>ttatggataaagcccgggtcattttgcccgcgataagtgatgcaataatgccaccaa<br>ccatggccatcaatgagttcattgcgaccagaccgctgatcccagaaatagcttgagc<br>cgacattacgttaaagccgaaccagcccacacaaagaatccaggcaccaagcgctaag<br>aatggaatgtttgatggtgcaaagttggtatgtttgcctgcgcgaatgcgccctttac<br>gcatacctaagaaaatgacggcaacgagagcaatccagccacccactccgtggacaac<br>gacggagcctgcgaagtcatgaaagccaaagccaaagcttgcttcaaaccaatcttgg<br>acaccaaaattaccgttccaaatgatgccttcaaacagcggataaacgatacctaccg<br>tgaaaaatgtcgcaatgaggataggataaaaacgcgcacgttcggcaattcccccaga<br>gacaatcgctggtatagcggcagcaaacgttagcaagaaaaagaatttaacaagctca<br>tatccattgccttgagacagtgtctgcgcatcggcaaagaaagtaccgccatacgcga<br>tccaatagccaataaagaagtaggctatggcggaaacaccaaaatcggccaaaatttt<br>aaccaacgcatttacttggttcttttgacgtacggtccctacctcaagaaaagcaaaa<br>cctgaatgcatcagaaataccatgatggcgccaagcaataaaaataaggtgtcggaac<br>tttgggtcagggtttgtacagctccgtatacctgacttacagtttgagtcattctcgt<br>tacactccatgcagcaatataaataatcgcaccaagcaggtgcttaaaaattatcaat<br>tcacaaaaatggtgaatttattatgtttgtacactgcttaggcaaaatgtgttccaga<br>ctgtttaaactagcttgcattgttatgcggtaaaacattaagcctatgaatattattg<br>atatatataaattttagtgagttttaagtgaaaggtttggtc |
| ***loxPsym* VIII** | ggcatgtgtttccagtctttcaacttgtttcctcaccttacggtattagaaaactgtg<br>ttttaccacagacttctaacttaggtactaacgaagaagtcgccattgagaaagcgtt<br>agaactgcttaataaggttaagatcaaagagcaagcgaacaaatatccaagccagctc<br>tctggcggccagcaacaacgcgttgcgatcgctcgtgcgttatgtatgaacccggaag<br>tgatgctgtttgatgagccgacttcagcacttgacccagagatgatctctgaagtgtt<br>ggatgtcatgaccgacttggccagtgaaggtatgaccatgctttgcgtgactcatgaa<br>atgggtttcgctcgtaaagtcgcagaccgcgttatctttatggatcaaggacagatcg<br>tagaagaaaataatccagaagaatttttctcagcaccaaaatctccacgcactcgcga<br>atttttaaatcagctgattcaccactaaggaagtatatgtttaatgagtattccatcg<br>ccattcatggtggtgctggtgctatgtctaaagggaaactaaccccctgagcaagagca<br>agaaatcgaagacattttaaaaggaattattgaagctggcttagctagtctccaagat<br>ggcgcaaatgcattagatgtggtgcaggcagcggtaaatcaacttgaagattgtacct<br>ggttcaatgctggcaaaggcgcggtattcaatcatgaaggaaaacatgagctagatgc<br>gtcgataatgtgcggtaaatctcttgatgcaggcgctgtatcgggcatacgctttagt<br>ccaaatccaatcgatgtcgcgcgtgcggtaatggataaatcgccacatgtttatttag<br>gtggtgaaggagctgagcagtttgtaagagatctaggtttagcacaggttgaaaatac<br>gtatttctctacggagatgcgttatcagcaactccagtctgcgttaaaagcccaagaa<br>gttatcctggagccaaccgataacgactacaaatacggtaccgttggtgctgtggtaa<br>aagacaaacatggtaatctcgcggctgctacatcaacaggcggcataactaataaaca<br>gtatggccgaatcggagactctcctgtaattgggtcgggcacgtacgcaaacaaccag<br>acctgtgcagtatctgcgactgggcacggtgagcacttttttgagacatgtcgttggtc<br>acaatatttcatctcgcatgtcattagccagtgagagcttagtacaggcggctaaaca<br>tgtggtatttgacgatttgccaagtacgggtggaagtggcggggttattgctgtcgat<br>agccagggtaatttgacccttccatttaacaccgaaggtatgtatcgcggttggggcg<br>gctcaaacgagcctgctcagtcaaaaatctatgaataacgctctgattatttattgtt<br>attaaccttaagagtaatcgtttcaaatgagaaataacttcgtataatgtacattata<br>cgaagttattgtgccctacaaccgaagtacaccgtagggcacatgttgtggagctggt<br>aagggtttagtaaatatatcgtcgaagttcgtgcgggtagacgcgcattccatttgag<br>gtgttttcgtaaacggattgttcttcgatataagaccagacataaggctctgtacttg |

| | |
|---|---|
| | tttgagcggcgagttgtgagatcaccgtatttacatcgtgaccagagtaatctttggg<br>atgaaccaacatttccttgatagctgacaacataataattccttttaatgtaattaaa<br>ttaccatttggtatgattaaaacgtagtttggtttttaatgtgatcaagtttggatt<br>ttgttttaatttaaaattcattttgtaagttaggaacgtagtttgaggcgcgacctc<br>actcacgttgcttaaggcactcaattttcattgactagtcatccagtgatgcgtggt<br>gagttctgtatttgtggtgtgataaaagaaaacggcgcaggtatcttgcgccgttttg<br>gtaattagctctttgagcttttaggtctaatcattattgagtgtagacgagttgacct<br>tctttgtaggtctttaacacttctatggtgttcatttttcaggttcgatggttagag<br>ggttggctgaaaggaccacaagatcggctagttttccaacttcaattgtccccttttg<br>gtcttcttcaagatattgatatgcggcgtttttagtgactgatgccaaggcttcatac<br>gcggttgcttttggtcttctcctagtgtctgaccagagcgtgtaactcggtttaccg<br>tgttgtagatgagtcgaataatatctgggtcaacaacgggagcatcgttatgagtggt<br>atagttaatgcctaagttacttgcagagcgggttggactgattcgaattgctcgctct<br>tcaccaaacacttcgtcgcgatgccaatctccccagaaaaaggtatgtgcgctaaaga<br>atgatgggatcatttgatatttgaccatcgcttctagttgatcatctcgagtggtttg<br>agaatgaatggaaaccggacgaattttgctaaagtcatcgctttccgttgaaattgcg<br>tcgagcgcggccaaatacacatcgacagacgcatcaccattagtgtgaacgatgaact<br>gagtgccttcgtctaggtaacttctcagcttagtttcgaggaactcgctggtcagggt<br>gggataacctacatagctggccgcttctccaactggtggatttaaatatggctgagtc<br>ataaacgcagttttgccttgtggtgaaccatcaaggacaattttagcaccgccaattc<br>taaagttatttgtatacgtcttgcttggtgcgtagccatccagtttcatcatgctgta<br>cgcatcccaagtcgggtaggcgacgatatctaagaacaaatcacctgacgctcctaaa<br>tcttcatagagtaaaactgattcatgactcgctgcaccttct |
| ***loxPsym* IX** | cgaagttttggcgattacagaccatgataccgttgacggtttagtgccagcgaaacag<br>tatgtagaagataatcagttgccgatcaaaattattaacggcatcgaaatctcaaccg<br>tttggcaaaataaagatatccatatcgtgggtctgaatattgaccctgataatccagc<br>attgaagacgcttattgaacaacaaaaacagcatcgaatcactcgttcagagctgatc<br>gcatcgcgtctgcaaaaagcaactcgtgaaggcgttctggaagaggtacagcagattg<br>caggtgatgcgccaattacacgtgcccatttcgcgaaatggttggtggacaatggctt<br>cgctaaaaacatgcaaatggtgttcaaaaagttccttacgcgtaacaaccctggttat<br>gtgccgccgaattggtgttcgatgaaagatgcgatagatgctattcacgctgctggtg<br>gccacgcggtgttagctcatccgggtcgatatcagttgacagcaaagtggattaagcg<br>tcttcttgcggcattctcggaagcaaatggtgacgcgatggaagtcgctcagccacaa<br>caagcacaacaagaaagacgcaatctggcggattatgctatacaatacaaactattag<br>cgtcccaaggtagcgactttcactatccgtctccttggatggagttgggtcgaaacct<br>ctggttgccgtcaggcgtagaaccagtttggaaagattggggcattgacccttcgttg<br>gatcgcaacgaagtccacgctccataaggatatagagccgagaggctcgataatgagg<br>aattacaatgagccagtttttttacgttcatccagaaacccacaggctcgcttgatt<br>aatcaagcagtagcaatcattcgtaacggcggtgtcgtggtttacccaaccgattcgg<br>gttacgcacttggctgtcagctggaaacaaacaggcgttagaacgtatctgtcaaat<br>tcgtcgcttggacgataagcacaacttcacgttgttatgtcgtgatttatccgagatt<br>tcgctgtacgcacgcgtagacaacggcgcatttcgattactgaaaaacaatacaccgg<br>ggccttacacattcatcttcaaaggtacaaaagaagtgcctcgtcgtttaatgaatgc<br>aaagcgtaaaaccataggtatccgtgttcctgataaccaaatcgcgcttgatcttttg<br>gaagcattgggtgagccattgatgtctacgtcgttgattttacctaacagtgacgtag<br>ctgaatctgatccagaagatatccgtgacaagttagaacatgcggttgacgtaatact<br>taacggcggttatttgggtgaacaaccgtcaacggttatcgactttagtgaaggtgac<br>cctgtcgtggttcgtcttggatcgggtgaccctgcgccatttgaataagtgtggtgct<br>agcttgatgtgataaaggaggctgagatgcctcataacttcgtataatgtacattata<br>cgaagttatctttttgtatctggcgtatcgattcgaaaacgaagtaaattctttaag<br>cgaactcgaaaagatcgatgcctttgtgatccgcataatcgatgtaactcgatgtgac |

| | |
|---|---|
| | gccattaagtgtcacttcaacattgtactcatcgctggagaataccgcccagccacac<br>tcattgtatgaaaggtcaagatcttctgctgagtcaagaaacgtatcgcgaatcccat<br>ccccgttagtatctcttccaaagtcgaaaaccaacttatccgagaaaatactgaaatc<br>cagtacttctgctatcacttcgatttcattagaagaaacaggttgatcaggaacacca<br>atcgccattctgttggtgaacacgaatgaatctgcgccaaagccaccacttaatacgg<br>actcaccatcaccgataacaatggtgtcattgccccaaccgccactaataaaattgtt<br>gcctgaataatcgaaaatgatgtcgtctccgaatccgccgtacagcgagtcatttcct<br>aaaccaccagacagataatcattaccgaagccaccagagatgaagtcgtttccccgcc<br>cgccgaacaagctgtcttcaccccagccaccgaaaacataatcgttcccgtcgaatcc<br>gacgaaattatctttaccgccaaaaccgaagaagatatcggttgtgtcggtgcctttg<br>gtaccagtcatattgtcatcgtcgcaggtgccaaatatccaattgagagtaagcgctg<br>tgtttgacatttcattgtcctttttatctgattatcttagcgggcaagtcagtaagtt<br>gttgttgtaacaaccgtgataattttatcagctgaagttggaacgcagcgggcgagag<br>agttctaaattagtacggttcttcgccagaatattgatggccagtaggaaaaaagtgg<br>tatatcggagttcaatcaggttacaagcagaaggcggagaatagagaggatttctctt<br>acattaaaccagaaagttacgtataatggccgactgaaaaaatcccgtgaagatgggt<br>aattgcataatacagaccaatcctggtctagacgtctgtgaagacgacacataggtag<br>ataaatgagcgaaaagttacaaaaagtattagcacgtgctggtcacggttctcgtcgt<br>gagattgaatcgttaattaaatctggtcgcgtaagcgtgaacggtaatgttgctaagc<br>ttggtgaaagacttgaagatgagaacagcgtggttcgtatcgatggtcacattgtatc<br>tgcgaagatacaagaagaagtgatctgtcgtgtactagcgtactacaagcctgaaggt<br>gaactgtgtactcgtcatgatccggaaggtcgccgtactgtttttgatcgcctgccaa<br>agattcgtggctcgcgttggatttcggtaggtcgtctggacgcgaacacatcaggtct<br>gttgcttttcactacggatggtgaattggcaaaccgcctaat |
| ***loxPsym* X** | agttagtttgccgtttacttctggaagtacaacgcctacagctttagcagcaccagtt<br>gaagatgggatgatgttctgagaagcaccacgaccaccgcgccagtctttagcagaag<br>gaccgtcaacagtttttttgagttgctgttgtagcgtgaactgtagtcataagaccaga<br>ttcgataccgaacttgtcgttaagaactttagcgataggtgctagacagttagtagta<br>caagaagcgttagaaacgatgtcttgaccagcgtaagtgtcgaagtttacgccgttaa<br>cgaacattggagttgcgtctttagaaggaccagttagtacaactttttcgcgccagc<br>agtgatgtgcttacgtgctgtctcgtcagttaggaaaagaccagttgcttcagcaaca<br>acgtcaacaccgatttcgtcccatttaggtcttcagggttgcgctctgcagttacac<br>gtacagttttgccgttaacgattaggttaccgccttcaacttcaacagtaccgttgaa<br>acggccgtgagttgagtcgtacttaagcatgtatgccatgtaatctacgtcgataagg<br>tcgttgatacctacaacttcgatgtcgttacgctcttgcgctgcacggaaaacgaaac<br>ggccgatacggccaaaaccgttaatacctactttgatagtcattatagttgctccaca<br>acttaatttctgattaaagataactggtagtaaaattacagaatccagtaaacatctg<br>caacagataatcggacttaacttgtttaaagtcaaaaaaaagcgacgctttttttaac<br>aattcttcgcaaatgttcgtttttttaaactttttactgcttttactctagctcagttg<br>gcgtaatataacctgcgagatgagtgctcgtatcaaattaattacacatggtgtcctg<br>gtgagaaagtatgtgctacaactgttacgagaatcaagtttttttgctaaaaaagtcat<br>tgcgaaaaacaggaaataaggatctaaaattagtggaacaaagcggaggaaaagtgac<br>gaagtcctatgaatattggcgtgagcgtctatctgatgaagagtttcgtgtttgtaga<br>gagcagggaactgagccaccgttcagtggtaagttattgcataataaggaaactggta<br>tatatagctgtacctgctgtaatgcccctctttttgtctctgataacaagtacgattc<br>aggatgtggatggccgagttttgatgcgccagttaatgaccaagcgatacgctattta<br>gatgatctgagtcacggaatggtacgtactgaaattcgctgtgcgacatgtgatagtc<br>atctggggcacgtttttgaagacggaccacaaacaacaggcgagcgttactgtgttaa<br>ttcagtgtcgttaattttcaacaaatctgaagattagtaacaaaattgcagttaagag<br>tgtttcatcctttcagaactgtatcggtcaacaataacttcgtataatgtacattata<br>cgaagttattataaacactgaaaacttgaatgaataaaaaactcccgactagcgggag |

| | |
|---|---|
| | ttttttatatattttcgggaattaaatcgggggaataacttccctcttcaatattat<br>ttgcgatcgactttgccaggtcgtccagttttgatattggtttggatcattgaaatc<br>gaacattctctgcagttgcttttcatctgctactgacacaccaaatcttttggctacc<br>attgcccaaaggtaagcacgtttttctccccttgtccctgatgaatactcgccaatg<br>atttgagtatggcagggtttaggttgtcttcatttgacaaacttaacgcattatttaa<br>caagataagcgttttaggggtgtctctgttgacgtagaaggtagctaaggcgtactgc<br>atttccgcagtgtttagtgtcggtgagccttccatctgtaaaaatcggtgctgagcct<br>gtgtgtctccggtctgtgaccaaaggaaatataaggtctctggcttttccgataccttt<br>taactcgctaacgatgcgttcaagctcgtcgatgctgtgcaccagcgcattaaagcgt<br>tgtgcttttagctcgctttggtcaattggagcgatctgggaggctaactccaagcact<br>ttctgtattcagcgacgacgccgtattcttctatctgtttttcttcgcttgggtgttt<br>gaactgctcataacgatgccaaattaagttggtacgcgggatgcggcattgtccatcg<br>tcgatgttgagcttttcgcagcgtagttctgggttatttttacacagttgttcagtag<br>atatgttgcgttcaaagcaacctgtgatggatagagtggcgaggcttgccagcaacca<br>gtatgttgttttcatggttttgcctgtgatccgttacaccaattcattttcgtgttct<br>tgactccatttcttaacccgctatgttctgaacagaatgtaataagaaggtacattat<br>ggacacagaacaattattaaaagcgatgacaccagaagtatacgatcgcttggtatat<br>gcagtagagacgggaagatggccagagggtacaacactttctagagagcagcgagatt<br>cgtgtatgcaagctgtgatgttgtatcaatcaaaacataatacagacgctcaacatat<br>gaccgttgctgccggtggtgaaatcagttttaagtctaagcaagaattgaaaaagcaa<br>ttcagtggtgatgagagcgatatcgttcgagtaaacccaaacttcgattgatacgttt<br>aacgcgcaaaaagcccaatgtaatcattgggctttttaggttttaagtgctgagaatt<br>gagtcttacaggctcatttcaccacgtaatacttgctgcatttttttcttttacttctt<br>cgtaagacaggttctggcttagaagataatgcagtttggctagtgcagcttcaggcgt<br>catgtcaaatccactgatcactccagcatcagccagcgcaca |
| ***loxPsym*** **XI** | gaacctactttgtcactgtcactgcagcaggcgagcctaatactgttgcaatagattc<br>tgctcctgtggatttgatgaacggtgtggtctatcaagtcgtagcgatagacgatggc<br>aatagtggtggtttcaatctgttagttaatgacatcacagactaggacggagtgggag<br>gggagactcctccctgcgataaggccgatatgatgagtgatagccgacaaaaactgg<br>gacggcatgaatggaacgcgtatatggacaaagtcaaagcacgagatcaagacgcctt<br>tgcttttgtattccaattttatgctcccaaattaaaacagttcgcttataagcacgta<br>gggaacgagcaagtcgcaatggaaatggtacaagaaaccatggccacggtttggcaca<br>aagctcacttgtacgatggaaagaagagtgcgttatctacttggatatacaccatcat<br>ccgcaacttgtgctttgacttattacgaaagcaaaagggcaaagagctacacattcat<br>gcggatgacatctggccttcagaatattatccaccagatctcgtcgatcactactctc<br>ccgagcaagacatgctgaaagagcaggtggtgaaattttagatacattgccgaaaaa<br>tcaaagagacgtgttgcaagcggtatatcttgaagagttgccgcatcagcaagtggca<br>gaactgtttgatatcccacttggcaccgttaaatctcgtctaagacttgcggtagaga<br>aattaagacattcaatgcatacggagcaactatgaacaatcatccagataacaaattg<br>ctagaagcttatgctttgggcagcattgatgcggtttctggtcttgtggtcgcgacgc<br>acttagaaacatgctccaagtgccgagactacgtcaatcgggtagaagcagagcaagc<br>gaatgtcgtcagtaacgttccttgtgtttacgtgcctgaatttgatgacatgtttaat<br>gccattgttacggcccctcccatgcgagacagtgtcattattagagactcagcaaaag<br>tgtccgtggcaggcaaaagctttgagttacccaaaactctgagccgtttctctgattt<br>ggtcggctcttggagaagttatggcggcaaagtatatagtgctcaaattgatttgggc<br>gaagatgccagagtaaacctcatgtatatcagtgaagatgtgcagattcctcaacata<br>cgcacaaaggaatagagtcaaccttggtcttacatggcggatttagcgatgaggatgg<br>tgagtacgaagagggcgacctgatgattcgtgatgcgtctgttaagcacagcccgtat<br>acgaaagccggggaagattgtttgtgtcttactgtcttaactgagccaatgatattta<br>ctcagggtgtggcaagaatcttcaatatgtttggtaaggggctttacccttaaactcc<br>agcagtttttcacaatgctgtttttaaaagaacataacttcgtataatgtacattata |

| | |
|---|---|
| | cgaagttatcaccgttttgcggtggttctttggtttctagtgatttagtgtttatggc<br>gtgcttttctagcaggtcatcaaagtagtggcgtagtgaatacagcatgatcaaact<br>tggtatcaccataaccgctgtaagaatgaagaaggtactccagtcattaaggtaatct<br>actagctcaccactgaatgaggccagagtggttcgtccaaagttgcctaaagacgcaa<br>gcaatgcgtattgtgttgctgaaaacgcttgtccggtcaataacgttaagaaagagac<br>aaacgccacggttgagaatgctgtagtaaagttatcgacaatgatggtagctaagtac<br>aagtgttcattaggaccgactgaagcaatccaagcaaacattagattacttgccgaca<br>tggcaatgccaccgatcatcaggccgcgaaccaagccaaacctcacattgaacatact<br>gcctactaaggtaaagaatattgtcgcgccccagccgattagcttggagtagtagcca<br>atttgctcattactaaagccgatatctttatagaatgcaatcgacatccggcctaaga<br>aggcttcgccaattttgaacaagaatacgaagagtaataaggttatcgcaacgcgcac<br>cccatttcggttgaagaaatccaggaaaggctcaacgacggttactgtaaaccacgca<br>acgattctagaaccaaccacttctttgtgacgttgttctgcttcagactgtaactgct<br>cacgttgagttttaggttctccgactaagagtgtaaacaccattaaaacaacaacgat<br>aaatgccattccgtaataaacgccattccagccgatgctatcagcattaataaacgct<br>aaatagccaggcagggagtagccagtccaccatccgatgactgccattgccgacgctt<br>gtggcagtttagaggcttcagattttgggaaggtatcaatacgaaaagcgtcgatggc<br>gatatcttgtgttgccgatgccgtcgcaatacacagggctaaaactgacgcaaaagcc<br>agattttgtgccgggtttacatctgcaataagaatggtggcgacaagtacgatgcttt<br>gacaaaagaaaatccagctgcgtcgctgcccaagaagacggtgcaacagtgggagttt<br>tagccggtctaccagtggagcccagaggaagttaatcgcatacacggcaaagacgcta<br>ccaaaataaccaatcgcagaaagcgttaagcctgcatctttaagccagccagacatgt<br>tagagccgatcaatacccatgggaagccgcttgagcatcccagcataaagacccataa<br>caaacgtttatcgaggtaactttttaccgtttccaaccaagtgatggaaggtaatgaa<br>gacatagattaatccaataaaatgacgccgttgaaagagaaaacccttcgaatattca<br>aagggtttatgatggctatttgagtaatgttgctttcgtgat |
| ***loxPsym* XII** | gctggaactgaccaaggagttgggcggcttaggtaaagtaggttgttcacttggtatg<br>aagcatcgtgaagataacctgaatcacggctaccagcattactcacaagaagtgatgg<br>aacaggaagcgttagcatcggttgaaaagcaaaagcaagcagagctaagtgacacgat<br>gtcttttgatgacttcctggaagactatttttcttatttaaaacaataacagagtagt<br>aaggattgggtgggattgaatattaagcctctggttgtatcggtgttaggtagtgtgt<br>tgctcgccggatgtgcgacggcaccaccacagcagcaagataatctatgtgagatatt<br>ccgcgaaaaatctggctggtatgatgatgcgaaagatatggaagaagagtggggtact<br>ccgatccatgtcgcaatggcgattatcaagcaagagagcggataccgacacgacgcaa<br>aaccacccaaagattatgtgctgggctttatcccttggggtagagtcagtagcgcata<br>tggctacgctcaggcgcaagatcccgcctgggacgattttcaggattcgaccgggcaa<br>ggtggctcgcgtactaactttgatgacgctatgatgtttgtcggttggtatacccacg<br>aaacccgtcgccagcttggtatctctttatgggacccatataaccagtatcttgccta<br>tcatgaagggagaggtggttatcagcgcggaacctataaaaggaaacctagcttagtt<br>aaagtcgcgagacgtgtagaacaaaccgcgaagaattatggctggcagctgaaacaat<br>gtcgtcaggaactggaagacaacagcagctggttcttttaaccctcatagggataaac<br>agaaatcaaaccaataccgtaaggtgttgagattacaatcaggagagatgtaatgcct<br>ttactcgatagttttactgtggatcacactcgtatgaatgcacctgctgtgcgagtcg<br>cgaaaaccatgcaaaccccaaaaggggacaccattactgtcttcgatttacgctttac<br>tgcaccaaacaaagatatcctttcagagaaaggcatccatacactagagcacctttac<br>gctggctttatgcgtagccatctaaacggtgacagcgttgagatcattgatatttcac<br>caatgggatgtcgtactggattctacatgagcctgatcggtacgccttcagagcaaca<br>agtcgcggatgcttggattgcatcaatggaagatgtactgaaagtagaaagtcagaac<br>aaaatccctgagctaaatgaataccaatgtggcactgcggcaatgcactcactggatg<br>aagcaaagcaaattgcgaaaaacattctggacgcgggtgtgtcagtgaataaaaatga<br>tgagctagctctgcctgagtcgatgttaaaagaactgcgtatcgactaagacctgagt |

| | |
|---|---|
| | tgtcttatcaaatgatttaacgaaatgagcctgataacttcgtataatgtacattata<br>cgaagttatacaatgtcaggctctttttatgtgtctgtaaacagattacttttagtta<br>ctgcctttcttccgtttagggtagacgcgaactaacttaattcggttctcttcaatct<br>cgacaatttccatcggatgtccggacacttgcacgctcaggtgactttcgggaatgtc<br>ttctaagtgctctaagatcaagccgtttagtgtacgaggaccatcagtcggcagatgc<br>catttcagacctttattgatgtcacggatattggtgctaccttcaatcaagaagctgc<br>catccccttgtgggctgatttcatcagataagcttggtgagatcgaagtggtaaactc<br>accgatgatctcttccaggatatcttcaagcgttaccaagccaataatgtcaccatac<br>tcatcaacgataaggccaatgcgctgtttattgcgctggaacttcaacatctgcacgt<br>tcagaggcgtaccttcagggatgtagtacacctcatcagccgcacgcagtaatgtctc<br>tttggtgaactcattcttttccagcatcaggcggtaagcctcacgcaaacgcagcatg<br>ccgaccacttcatcgatttggtcgcggtaaaaaacgatacggccatgtggtgagtgcg<br>ttaactgacgaacgatagacttccagtcgtcgttgatgtcgataccagtaatttcatt<br>acgtggcaccataatatcgtttaccgtgacgtgctcaagatctaggatggacaccagc<br>atatcttggtggcgacgagggataaggccgccggcttcatttactactgtacgtaatt<br>cttctgagctgaggtgatcttccacagagtgatccgcctttaccccaagcagacgaat<br>aaagccattggtaatgaagtttacaagcagcacaagtggtgaaagcaacttcatcaga<br>atggtcaacagaatgctgctggtataagacacgcgctctggatacaaagcggcaatag<br>tttttggcgtcacttcggcaaacaccaaaaccaccatggtgagcgcaccagtcgcaat<br>cgccacacccatgtcaccgtagaggcgcatgccgataatagtcgcaattgcagaagcg<br>aggatgttcacaaggttattgccgataagaataagaccgatcaggcgatccggacggt<br>ccaataatttttcgacacgctttgctcccttatgcccatttttggctaagtgtttcaa<br>gcgataacggttcagtgacatcatccctgtttctgaaccggaaaagtaacctgatatg<br>acgataagacacgcgagtagcgcaaataagatacccgtagatatgtcgtccaaaacgt<br>tgctgttccttgtaagtattgggttatttttaaaagctatgtataactgttaagtcat<br>agatttacggtactgtatttgatttaactttgaatataaagggcttgtggacgcctta<br>attcaaaatgatctcttttacaaagcggctgccgaaataagc |
| ***loxPsym* XIII** | tcaaaccctgcaaatttgacttacctcggcactactccccctgaataattcgtattgg<br>aattgcggttccacaaattatctacctgggtagtgctcctcttccacataaagctcaa<br>acattgaacggttttggcgtcacaaagtcaatgaatatttcaactattccaagatttt<br>ttattagtgaaattgtgacagtggtttgaactcaatacctacggctaattgccgagca<br>ctataaaatccattagtgtgagttctttgatgcccgagacttattcaaaattctaaaa<br>actattgtccgcgctcgggttaagtaacgtcaaacaaggagctcacatgccgatccaa<br>agctttattccccctcaccgtaccctaatgggaccaggtccttcagatatctatccgc<br>aagtgctgcaagcgcttagccgcccaacggttggtcacttggaccctctgtttatcgc<br>aatgatggatgagctgaaacaattgctaaagtacgcgttccagacagaaaacgacttt<br>actattgccgtttctgctccaggtagtgcgggtatggaggcatgttttgttaacctta<br>tcgagaaaggtgacaaagttatcgtatgccgaaatggtgtttttggtgagcgcatgcg<br>tgaaaacgttgttcgtgctggtggtgaagttgtcttggttgatgacgagtggggcgcg<br>ccagtctctgtcgataaagtcgaacaagcactgcagcagcatcctgacacaaaattt<br>tagcatttgttcacgcggaaacatcgacaggtgcggtcagcgacgctcaagctttggg<br>taagttagccaagcaacacaatgcgctctctattgtcgacgcagttacgtcattaggt<br>ggggtaccgttaaaagtagatgaatggcaactggatgcagtgtactcaggtagccaaa<br>aatgtttgtcgtgcgtacctggcttgtctccgttgacattctcccaggctgccattga<br>taagattcaatcgagaacaacaccagtacaaagctggttccttgatcaaagcttagtg<br>cttggttactggagtggagagggtaagcgcagctaccatcatacagcaccagtgaata<br>gcctgtatgctttgcatgaagcgttattggttctgcaaaatgaaggtttggaaaatgc<br>ttgggctcgtcaccaattgatgcatgaaaaactaaaagaaggtctacaaaagctcggt<br>tttgagtttgttgttgaagaagcgcatcgcctacctcaattaaatgctatttatgtac<br>cggaaggtattgatgaagcgaaggttcgagctcacttgctggaaacctataatctgga<br>aattggtgcaggtttgggtgccttggcaggcaaagcgtggcgtatcggcttgatgggc |

| | |
|---|---|
| | tacgcagctcgtccagaaaacgtcgcgctatgtttgaaagcgctggaagagtcgctga<br>cagagtagtgacacctaaagtaaaaaagcgaagataacttcgtataatgtacattata<br>cgaagttatgttatccttcgctttttttcgtttatttactgctcgtctgacagcttagg<br>tgaagcagattgagcttgtgatggtggttggtaattcactttactgaaatctacccga<br>ggaaataagaactgcgcctctgcacgaatttgctcggctttacttccatcgtctagtc<br>cctggtaagccaggatcaaattgttatagaaggccgggcgtggcttatctttaatgat<br>attcagtgaccaatcaatgtaaggctgaatcaagcttggatcttgtgtttgtaacccg<br>atagttagaaaggtgctgtacacatcccagtcgaatctatctttccaaaccacagggt<br>tactgacttggctaagaatatctgggtttgtcggacgtgtggtttcaaatttcgttag<br>tatgtagttggtatgtaaggcactcaccatgtagaacgtaaacacggctggcacagcc<br>aagcttacgatgcgcagcaagcttttcgtgattttttgtaaagccgccttggcgataac<br>gcgcaactcgctgatccacccaatatattaaaatgataaagattacccaatgcactaa<br>cgagtgataaaacgggtattcgagctgcgagtggagaacaatgggtataaacaatgcc<br>agtaaggctaaacgtgtcccgcgcttggcctgatagattctgtgcagcactaatgtca<br>tggccaagaagataccgataattggcagtaacccaccttctacggcccagtacaataa<br>ttcattgtgcgggtgatccattgcaggcaatcccgcaggatatttctcatttaacgaa<br>tgttggcgagcggtataaatcatgtactccgactcaaactttccgtaaccatagccgg<br>tgaacggtttctcaatcaccatatctaccgtttgcgggaaagtgtaagcgcgagccga<br>ttccatgttgactttctcggcggctaacccgttaccttgtgggaaagcgatttgcatg<br>accactactgataacaacagcccggctgcgagtgaaatcacccaatgcgtaaagcgcc<br>gcttggtattgtagcggtacatgtacgggataatcagaaggaccgcaaacagagaagc<br>cagccaacctgttcgtgaagcaagcgcaacgatcaaaggtatcgttaaaacgggtacc<br>gcataaagtaagtagacatcactgagcttacgatcgtacttgtatggctgacgcgcca<br>gtaagtaactggcaatgactaagccagttgctagaaagctagccatgacattgggttg<br>ttgaaagataccgtatggacggttagctacggtgttgtatccgaaaggattatctggt<br>tttaagtagagatattgcgtcagaccgaaaatagcctctatcaccaccgctaacacga<br>taaaccacaagattcgttgacgatgcttattgctgaaatggaattgctgaagtacaac<br>gaaaaacagtaaaccactccacaggccaatgagtttattcgc |
| ***loxPsym*** **XIV** | taagaagatcggtgttttaacaagtggcggtgacgcacctggtatgaacgctgcagta<br>cgcggcgtggtacgcacagcgctttctgagggattggaagtgtttggtgtgtacgatg<br>gctacctaggtctttatgaaggccgcatcgagaagctagatcgttctagtgtttctga<br>tgttattaacaaaggtggtacgttcctgggttctgcacgtttccctgagttcaaggaa<br>gtggcagtacgcgaaaaagctattgagaatctgaagaaacacggtatcgacgcactgg<br>tcgtgattggtggtgacggttcttacatgggtgcaaaaaaactgactgaaatgggtta<br>cccatgtatcggtctgccaggcactatcgataacgatattgcgggtacggattacact<br>atcggttacctaactgcactaaacacagttattgatgcaatcgaccgtctacgtgaca<br>cttcttcatctcaccaacgtatttcaattgtagaaatcatgggccgtcactgtggtga<br>cctgactctgatgtctgctatcgctggtggttgtgagtacatcatcacaccagaaacg<br>ggtcttgataaagagaagctgatcggcaacattcaagacggtatcgcaaaaggtaaga<br>aacacgcaatcattgctcttactgaactgatgatggatgcaaacgagcttgcacgtga<br>tattgaaacggcaacaggtcgtgaaactcgtgcgacagtactaggtcatatccaacgt<br>ggtggtcgtcctactgcatttgaccgtgttcttgcatctcgcatgggtaactacgcgg<br>tccatcttctgatggaaggtcacggtggccgttgtgttggtatcgtgaaagagcagct<br>tgttcatcacgatattattgatgcgatcgaaaacatgaagcgccctgtacgtaatgac<br>ttgtacaaggttgcagaagagctgttctaagtcgaccgctcttagctaaaataaaaaa<br>ccgcgttgatacgcataacttcgtataatgtacattatacgaagttatggtttttttg<br>tctctgtttttctaaagctaggggaggacagggttcatgaagaggaaatgctgtgttc<br>taccagcattaactttcgttcgcgaagcgcaaacatgtaaccaagcaacccacctaat<br>atattacccaccgcaataccgataaatagcccttcaacatcgtacaggtagctgccaa<br>tccaagctgcaggcagcatgaataagaacaagcgcataaagctccattggaaagcgcg<br>taatggcttgtgcatcgcattcagcccgctcactaatatcatcacaataccctggaaa |

| | |
|---|---|
| | ccgtagctgaacggcacaacaaggagataatgccaaagcagattttaactgcttctt<br>cttgtgagaaaagagccgccagagggatgctcaaaggcaccattgcaaggaagataag<br>cccctgaaacacaactgcgaagcgcatgctgaggaataatcctgcgaatgcccgctgt<br>gggttattagcgccgagattttgtgccatgaacggcgttaaagctgaggtcagtgaca<br>tcaagaccagaatcagtatcgactcgatgcgctgtgcggcgccataagcggcaacggc<br>tgcggttccgtggcttgacagcagcatcatcaaaatagcgccagacagcggcgtcatt<br>gcggtagagaaggccgctggcgtaccgattttcaatgtctgttgccaatcttgtttca<br>gatgtaaccactgcggcttagccagaagtttctcccgtttaactagtaggtatgacga<br>gccgactaaagcgcctaaccaactaaaagcactcgcaattgctgcgccctggattcca<br>agttcaggaaaaggcccaaagccgaagatcagtagcggatccagcattccgttgatca<br>accctgccagcatcatgattttagctggggttttcgtatcacccgtcgcacggattgc<br>actgttgccagccattgggatcaccagtaatggaatggtgagataccacacttgcatg<br>tact |

**Table S2. Primers used in this study.**

| Name | Sequence 5' - 3' |
|---|---|
| 1F | agaagaagcgggttctccagtcg |
| 1R | agacttaagtacctaggcccttccatcg |
| 2F | ggcggagttggttcgcaataagc |
| 2R | gaaaaggctggctgacatggctg |
| 3F | tagcgaaagctgtgcgaagtttctg |
| 3R | cgcacttgcgacccttttgttaagaac |
| 4F | agcggcattgaaagtgttgctattcag |
| 4R | aagtaacgtttgggacgcaactgc |
| 5F | gctttgtgatgtggttggtgagtcg |
| 5R | ctcgcctgctcatcaaccattgaag |
| 6F | cgaagattaacaaaatgcaaaagagcg |
| 6R | ggcgtaatgatatctctggtgaaggtcg |
| 7F | cacaacatgtgccctacggtgtac |
| 7R | ggaaattgctagaacgtattgaaacgctcg |
| 8F | catcgattatgcggatcacaaaggca |
| 8R | ctcaaacgagcctgctcagtcaa |
| 9F | gacctggcaaagtcgatcgcaaat |
| 9R | gtcgtggttcgtcttggatcgg |
| 10F | cactagaaaccaaagaaccaccgcaa |
| 10R | caggcgagcgttactgtgttaattcag |
| 11F | ctaccctaaacggaagaaaggcagtaact |
| 11R | ggctttacccttaaactccagcagtt |
| 12F | ccaaccaccatcacaagctcaatctg |
| 12R | ctgcgtatcgactaagacctgagttgt |
| 13F | gtgggttgcttggttacatgtttgc |
| 13R | cagctcgtccagaaaacgtcgc |
| 14F | acgaaggtggatagccttgtaagcg |
| 14R | gaagcgccctgtacgtaatgacttg |
| Δdns_specR_F | agagaacatagcgttgccttgg |
| Δdns_cmR_F | caagatgtggcgtgttacgg |
| Δdns_R | gtcacttaaagcgagtgtac |
